# Comprehensive profiling of transcription factors for reprogramming human astrocytes to neuronal cells through endogenous CRISPR-based gene activation

**DOI:** 10.1101/2025.10.11.681828

**Authors:** Samuel J. Reisman, Dahlia Halabi, Samantha E. Miller, Lingyun Song, Sara Geraghty, Nicholas Sangvai, Grayson Rice, Alexias Safi, Gregory E. Crawford, Charles A. Gersbach

**Affiliations:** Department of Cell Biology, Duke University, Durham, NC, USA; Center for Advanced Genomic Technologies, Duke University, Durham, NC, USA; Department of Biomedical Engineering, Duke University, Durham, NC, USA; Department of Biology, Duke University, Durham, NC, USA; Department of Pediatrics, Duke University Medical Center, Durham, NC, USA

## Abstract

Neuronal loss is a hallmark of neurodegeneration and brain injury. Direct reprogramming of astrocytes into neurons has emerged as a promising approach to restore lost neurons. Comprehensive mapping and characterization of candidate astrocyte-to-neuron reprogramming factors is an essential step to realizing the potential of this strategy. Here, we established a CRISPR activation (CRISPRa)-based approach for neuronal reprogramming of primary human astrocytes. We conducted high-throughput CRISPRa screens of all human genes encoding transcription factors (TFs) to identify novel and efficient reprogramming factors. scRNA-seq characterization of top hits revealed that single TFs reprogram primary human astrocytes into multiple neuronal subtypes with distinct cell type-specific gene signatures. We demonstrate that *INSM1* reprograms astrocytes to a glutamatergic neuron-like state and has broad neurogenic activity across different cell types and across human and mouse contexts. Finally, we conduct paired CRISPRa screens to identify cofactors that cooperate with *INSM1* to enhance neuronal reprogramming and subtype specification, and elucidate genomic mechanisms of interaction and downstream regulators.

## INTRODUCTION

The mammalian brain lacks sufficient regenerative capacity to restore neuronal populations lost to brain injury or neurodegeneration. As a result, cell and gene therapies that aim to regenerate neuronal populations have gained significant attention. Direct reprogramming of glial cells to a neuronal state has emerged as a particularly promising approach for generating new neurons^1–10^. Astrocytes are attractive candidates for neuronal reprogramming due to their shared lineage during neurodevelopment, abundance across brain regions, and latent proliferative potential which can become activated in response to injury or degeneration^11,12^. Direct reprogramming of astrocytes in situ circumvents major barriers associated with iPSC-based cell-therapies for neuroregeneration such as the difficulty of generating, expanding, differentiating, and transplanting patient-derived or allogeneic cells at scale. However, previous reprogramming approaches have demonstrated mixed efficacy and efficiency, highlighting the need for an expanded toolkit of technologies and reprogramming factors^13^. Moreover, neurons consist of many subtypes with distinct regional distributions, functional roles, and disease associations. For instance, Alzheimer’s disease and ischemic stroke result in the loss of glutamatergic neurons^14,15^, epilepsy (MTLE) and Huntington’s disease result in the loss of GABAergic neurons^4,7^, and Parkinson’s disease results in the loss of dopaminergic neurons in the substantia nigra^16^. Therefore, there is a critical need for high-throughput mapping of factors able to efficiently reprogram astrocytes to neurons and a comprehensive understanding of neuronal subtypes resulting from cell reprogramming.

Here, we address this need by developing a CRISPR-activation (CRISPRa)-based approach for astrocyte-to-neuron reprogramming and conducting high-throughput CRISPRa screens of all genes encoding human transcription factors (TFs) in primary human astrocytes. These screens and subsequent characterization with scRNA-seq reveal that CRISPRa of single TFs is sufficient to broadly modulate gene networks and drive reprogramming to multiple neuronal and glial cell states. In particular, we identify *INSM1* as a robust reprogramming factor with broad pro-glutamatergic neuron activity across cell types and both mouse and human species contexts. We further conduct paired screens to identify cofactors, most notably *IKZF1,* that cooperate with *INSM1* to enhance neuronal reprogramming efficiency and subtype specification. This approach can be utilized more widely for discovery of factors able to drive relevant phenotypes in other primary cell lineages. Overall, this work widely expands the list of potential factors for cell reprogramming-based neuroregenerative therapies and provides new insights into the TF interactions and gene regulatory networks (GRNs) that govern neural cell states.

## RESULTS

### Primary human astrocyte characterization

Validation and characterization of starting material is critical for cell reprogramming experiments. Thus, we first performed RNA-seq of primary human astrocyte cultures and examined expression of cell lineage markers to determine astrocyte subtype and culture purity (**Figure 1A**). Profiled cells expressed high levels of the canonical astrocyte marker GFAP and markers of both upper layer (BSG, ITM2B) and deep layer astrocytes (ID3, DKK3). We did not detect significant expression of contaminating neural cell types, including immature or mature neurons (DCX, MAP2, RBFOX3/NeuN), radial glia (FOXJ1), oligodendrocyte progenitor cells (OPCs) (PDGFRA), oligodendrocytes (PLP1), microglia (PTPRC), or other neural cell lineages. This was supported by immunofluorescent imaging where we also observed astrocyte morphology similar to primary human astrocytes isolated from immunopanning^17^ (**Figure 1B**). An important function of astrocytes in the CNS is to tune synaptic function and prevent neuronal excitotoxicity by uptaking extracellular glutamate^18^. We examined glutamate uptake dynamics to functionally validate primary human astrocyte cultures and observed robust depletion of supernatant glutamate. Further, to confirm this activity was driven by glutamate transporters, we treated cells with 1mM 4-aminopyridine (4-AP), a potassium-channel blocker which stimulates glutamate release by reversing transporter activity^19^, and found that treatment resulted in effective restoration of supernatant glutamate levels (**Figure 1C**). Together, these data indicate a generally pure culture of functional cortical astrocytes.

**Figure 1.**
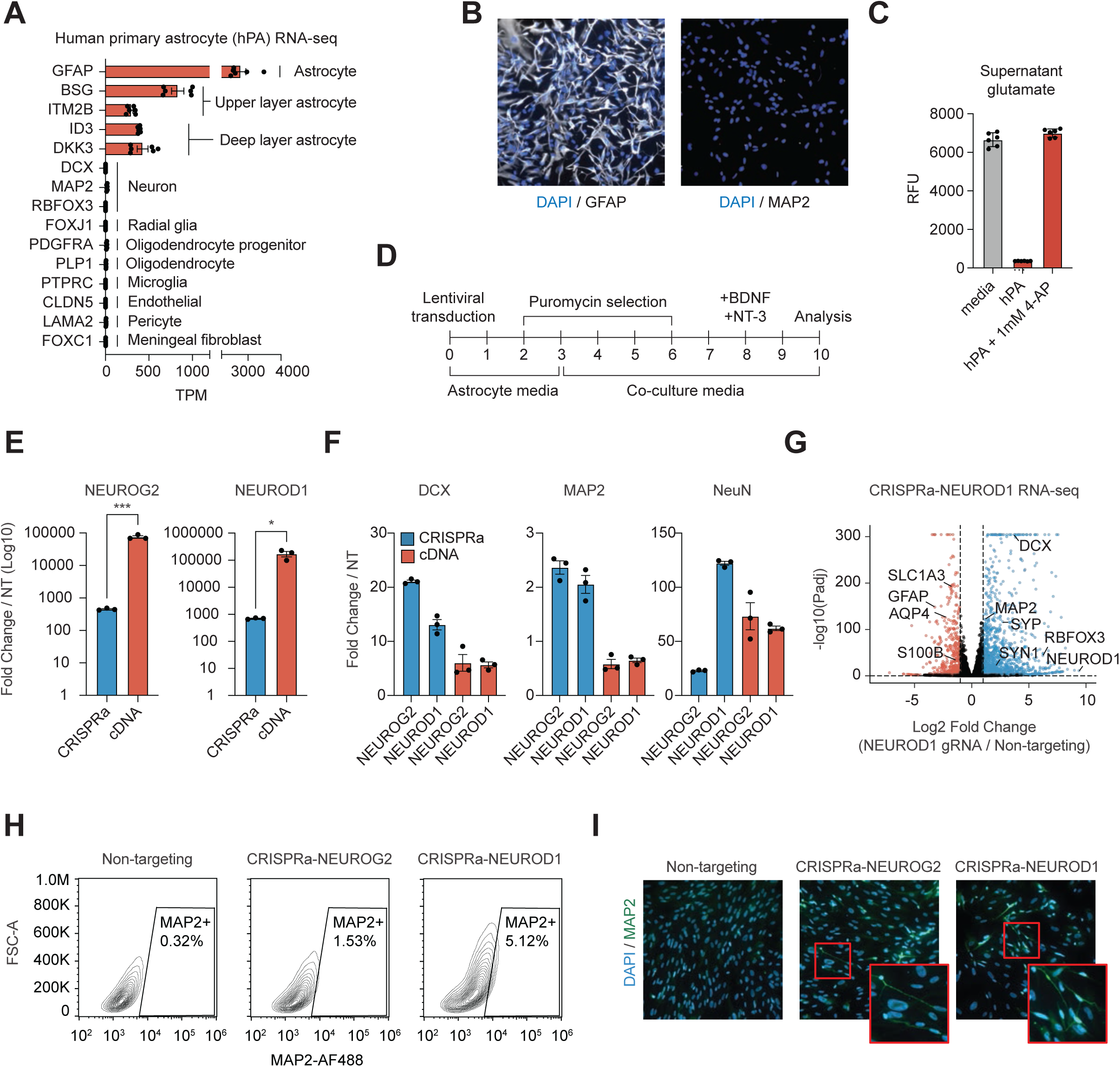
CRISPRa of known neurogenic transcription factors reprograms primary human astrocytes to neuron-like cells at low efficiency. A. TPMs of cell type lineage markers in primary human astrocytes (hPA) before treatment (RBFOX3 is the gene symbol of NeuN). B. Immunofluorescence staining of astrocytes assessing GFAP and MAP2 expression in primary human astrocytes before treatment. C. Functional validation of human primary astrocytes with glutamate uptake assay. *p < 0.05, ***p < 0.001 by global one-way ANOVA with Dunnett’s post hoc test comparing all groups to media; error bars represent SEM. To reverse transporter activity, 1mM 4-aminopyridine (4-AP) was added to cell culture media for 30 minutes. D. Schematic of reprogramming protocol. E. Comparison of RNA levels of NEUROG2 and NEUROD1 after CRISPRa with a promoter-targeting gRNA or cDNA overexpression with tet-ON system in constant presence of 2 ug/ml doxycycline. *p < 0.05, **p < 0.01, ***p < 0.001 by two-tailed t-test. F. Comparison of RNA levels of neuron marker genes DCX, MAP2, and NeuN as proxy for neuronal reprogramming. G. Differential expression (DE) analysis of RNA sequencing of ^VP64^dSpCas9^VP64^ + NEUROD1 gRNA (CRISPRa-NEUROD1) vs. ^VP64^dSpCas9^VP64^ + non-targeting gRNA, 10 days post-transduction. Canonical astrocyte marker genes (left) and neuron marker genes (right) are labeled. DE genes are determined using a Wald test by DESeq2 (padj <0.01). H. Flow analysis of intracellular MAP2 expression 10 days post-transduction. I. Immunofluorescence staining of reprogrammed cells to assess MAP2 expression 10 days after transduction. Insets highlight sparse neuronal morphology.

### CRISPRa of a single factor reprograms primary human astrocytes

Next, we tested whether CRISPRa of a single neurogenic TF would be sufficient to reprogram astrocytes into neurons. CRISPRa acts on the native genomic locus of targeted genes, allowing the retention of endogenous regulatory elements and the capability to upregulate multiple gene isoforms^20–23^. In contrast, cDNA overexpression is exogenously expressed, often at supraphysiological levels, and typically limited to a single isoform. Therefore, we hypothesized that CRISPRa would be effective in reprogramming cell state by more closely mimicking native gene regulatory networks. To test this hypothesis, we targeted known neurogenic bHLH transcription factors *NEUROD1* and *NEUROG2*, well-established drivers of neuronal differentiation and direct reprogramming^24–27^. We either overexpressed cDNA or delivered the CRISPR-based activator ^VP64^dSpCas9^VP64^ and a promoter-targeting gRNA^28,29^. Three days after lentiviral transduction (D3), media was changed to a basal co-culture media to support survival of reprogrammed cells, and on D10 we performed RNA-seq, flow cytometry, and immunofluorescent imaging to assess cell state (**Figure 1D**). As expected, cDNA delivery resulted in greater expression of the targeted TF than CRISPRa by multiple orders of magnitude (**Figure 1E**). However, CRISPRa led to higher expression of downstream neuronal marker genes DCX and MAP2, and CRISPRa-*NEUROD1* resulted in the highest expression of NeuN (**Figure 1F**). These results indicate that the more physiological levels of expression from endogenous gene activation support more effective coordination of the desired downstream gene regulatory networks.

We next assessed the transcriptome of reprogrammed cells. RNA-seq of CRISPRa-reprogrammed cells revealed extensive transcriptome remodeling, with CRISPRa-*NEUROD1* and CRISPRa-*NEUROG2* resulting in 6257 and 3440 differentially expressed (DE) genes respectively, including upregulation of canonical neuron markers and downregulation of canonical astrocyte markers (**Figure 1G**, **S1A-C**). Gene ontology analysis revealed CRISPRa reprogramming overwhelmingly increased expression of genes associated with neuronal processes (**Figure S1D,E**). However, flow cytometry and imaging of MAP2, a known neuronal fate marker^30^, revealed that the observed transcriptomic changes after targeting known neurogenic bHLH TFs were driven by a small subpopulation of cells, even though all cells received the CRISPRa construct (**Figure 1H,I**).

### High-throughput CRISPRa screening identifies novel astrocyte-to-neuron reprogramming factors

Encouraged by the potential of CRISPRa to induce transcriptomic changes but unsatisfied with reprogramming efficiency of known neurogenic TFs, we performed a high-throughput CRISPRa screen of all genes encoding annotated human TFs (the “TFome”) to find new and efficient factors for astrocyte-to-neuron reprogramming^31^. Primary astrocytes from three donors were transduced at low multiplicity of infection (MOI) with an all-in-one lentiviral library encoding ^VP64^dSpCas9^VP64^ and a single gRNA (**Figure 2A**). The gRNA library consisted of 6 gRNAs targeting the TSS of each TF gene and 5% non-targeting (NT) gRNA controls, for a total of 10,233 gRNAs across 1,627 TF genes (**Figure 2B**, **S2A**, **Supplementary Table 2**). After antibiotic selection and culture, we fixed, permeabilized, stained, and sorted cells into bins based on MAP2 expression, with high MAP2 expression serving as a proxy for cells reprogrammed to a neuron-like state.

**Figure 2:**
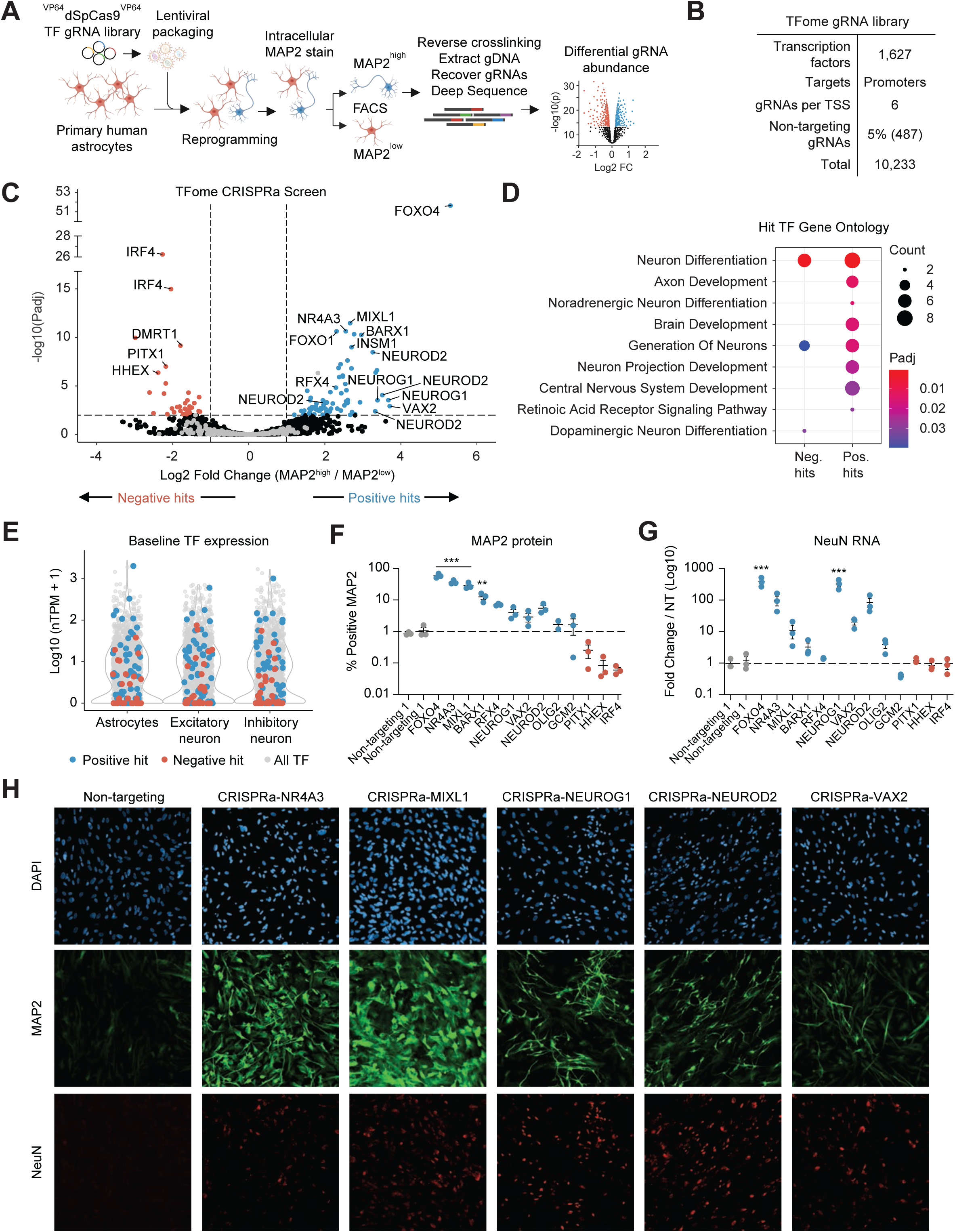
High-throughput TFome CRISPRa screen to identify novel TFs for astrocyte to neuron reprogramming. A. Schematic of CRISPRa screening with TFome gRNA library. B. Summary of TFome gRNA library. C. Significance (P_adj_) versus fold change in gRNA abundance between MAP2-high and MAP2-low populations in TFome CRISPRa screen. D. Selected enriched biological processes for positive and negative hits from the TFome screen. Statistical significance was determined using a two-tailed Fisher’s exact test followed by Benjamini–Hochberg correction. E. Expression level of hit TFs in astrocytes or neurons in the Human Protein Atlas Single Cell Type data. F. Validations of selected hit factors for MAP2 protein expression 10 days post-transduction. **p < 0.01, ***p < 0.001 by global one-way ANOVA with Dunnett’s post hoc test comparing all groups to non-targeting 1, gating set to 1% positive for non-targeting gRNAs; error bars represent SEM. G. Validations of selected hit factors for NeuN RNA levels 10 days post-transduction. *p < 0.05, ***p < 0.001 by global one-way ANOVA with Dunnett’s post hoc test comparing all groups to non-targeting 1. Error bars represent SEM. H. Validation of selected hit factors for NeuN, MAP2, and neuronal morphology by immunofluorescence staining.

In total, 70 gRNAs targeting 62 TFs were significantly enriched in the MAP2-high bin (p_adj_ < 0.01) and were deemed “positive hits” (**Figure 2C, S2B, Supplementary Table 2**). These positive hits included multiple gRNAs targeting known pro-neuronal bHLH TFs including NeuroD and NeuroG TFs. Positive hits also included novel TFs not previously tested for astrocyte-to-neuron reprogramming. Gene ontology analyses of the hit TFs revealed enrichment for neuron differentiation, axon development, brain development, and other neuron-related terms, indicating that the screen uncovered a range of TFs associated with neurogenesis (**Figure 2D**). The screen also revealed that the ability of a TF to reprogram astrocytes to neurons was not predictable by its expression level in neurons vs. astrocytes, supporting the value of unbiased high-throughput screening over candidate selection based on expression level (**Figure 2E, S2C**). For some hit TFs, only a single gRNA targeting that TF emerged as a significant hit. However, for many of these TFs, the other screened gRNAs were enriched in the MAP2-high bin but did not reach statistical significance. However, together they resulted in a significant change in p-value distribution compared to gRNAs targeting non-hit TFs (**Figure S2D**).

We next individually validated top gRNAs from the screen and assessed reprogramming efficiency. We transduced astrocytes from different donors than were used in the screen with gRNAs targeting ten strong positive hits (*FOXO4*, *NR4A3*, *MIXL1*, *BARX1*, *RFX4*, *NEUROG1*, *VAX2*, *NEUROD2*, *OLIG2*, *GCM2*) and three strong negative hits (*PITX1*, *HHEX*, *IRF4*). Positive hits increased MAP2 expression in individual validations, while negative hits decreased MAP2, confirming that the screen accurately detected changes in MAP2 expression across reprogramming efficiencies (**Figure 2F, S2E,F**). Additionally, the most efficient identified hit outperformed the efficiency of NeuroD1 in previous tests by an order of magnitude (58% vs. 5% converted). Most positive hits also upregulated expression of NeuN, a nuclear marker of post-mitotic neurons, indicating that activation of hit TFs resulted in upregulation of multiple neuronal genes rather than simply directly targeting MAP2 (**Figure 2G**). We used immunofluorescence imaging to evaluate neuronal morphology after reprogramming with a subset of our hit neurogenic TFs (**Figure 2H**). Reprogrammed cells displayed a range of neuron marker expression and morphologies, suggesting that the TFome screen uncovered hits with heterogeneous downstream effects.

### scRNA-seq characterization of candidate astrocyte-to-neuron reprogramming TFs

To comprehensively characterize the effects of candidate neurogenic TFs and profile the subtypes of reprogrammed cells, we conducted a follow-up perturb-seq screen. We cloned a sublibrary consisting of all significant gRNA hits from the TFome screen and non-targeting gRNA controls for a total of 119 gRNAs across 90 TFs (**Figure 3A, S3A, Supplementary Table 3**). We transduced astrocytes with this library at an MOI of ∼0.5 and profiled cells with single cell RNA-sequencing (scRNA-seq) with gRNA capture (**Figure 3B**). In total, we profiled ∼47,000 high quality cells across 2 donors (**Figure S3B,C**). Some cell clusters expressed astrocyte markers, largely on the right side of the resulting UMAP, while other clusters strongly expressed neuron markers, largely on the left (**Figure 3C**, **S3D,E**). To determine the cell types present in these clusters, we assessed expression of the gene sets that mark all neural cell types present in three scRNA-seq atlases of human development, which we called “Fan, “Zhong”, or “Descartes” based on the first author or atlas name^32–34^ (**Figure S3F**). We projected these expression scores onto our cell distribution (**Figure 3D**). Enrichments were concordant across all three atlases, with astrocyte signatures centered on clusters 1 and 3, glutamatergic/excitatory neuron signatures centered on cluster 15, and GABAergic/inhibitory neuron signatures centered on cluster 10. We therefore concluded that profiled cells displayed a spectrum of neural cell states including multiple neuronal subtypes.

**Figure 3:**
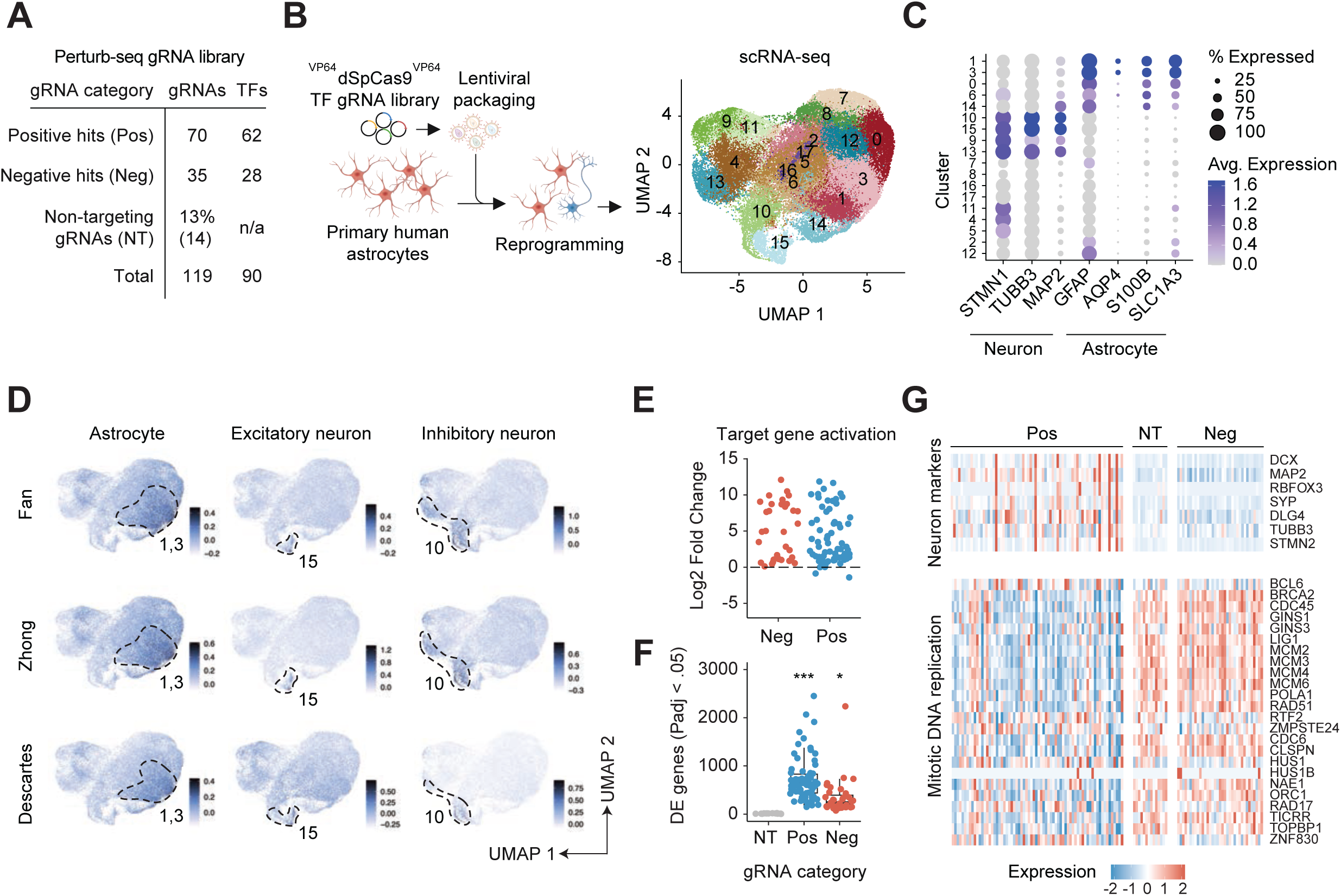
scRNA-seq validation of candidate astrocyte to neuron reprogramming TFs. A. Summary of TFome hit library. B. Schematic and resulting UMAP of Perturb-seq screen of TFome hits. C. Dot plot of expression of astrocyte and neuron marker genes in each UMAP cluster. D. Module scores of neural cell type gene signatures from scRNA-seq atlases overlaid on UMAP. E. Fold change in target gene expression for negative (n=35) and positive (n=70) hit gRNAs. F. Differentially expressed genes (DEGs, significant gRNA–gene links) were defined using a two-tailed MAST test with Bonferroni correction^95^. Significance of differences in number of DEG numbers were determined by a global one-way ANOVA with Dunnett’s post hoc test comparing all groups to NT, *p<0.05, ***p < 0.001. G. Expression level of neuron marker genes and genes associated with mitotic DNA replication by gRNA.

We then assessed the effect of each CRISPRa perturbation by comparing the transcriptomes of cells that received each gRNA to cells that received only a non-targeting gRNA. We first examined on-target TF activation and found that almost all gRNAs robustly activated their respective target gene (**Figure 3E, S4A**). We next examined MAP2 expression in the scRNA-seq data to cross-validate the FACS-based TFome screen. Almost all positive hits increased MAP2 expression (fold change > 0 vs. non-targeting gRNA) and the majority of negative hits decreased MAP2 expression (**Figure S4B**). Additionally, there were many other differentially expressed genes (DEGs) associated with each perturbation. Positive hits had the greatest number of associated DEGs (mean ∼654), while negative hits had fewer (mean ∼436), supporting that the MAP2-high bin in the FACS-based screen represents a genuine change in cell state (**Figure 3F**). The level of background noise in the DEG testing was low: on average, non-targeting gRNAs were associated with very few significant DEGs (mean ∼13). Robust DEG testing was enabled by high coverage of our gRNA library (mean ∼692 cells per gRNA). However, differences in gRNA representation did not lead to differences in number of DEGs (**Figure S4C**). Positive hit gRNAs, which had the most DEGs, were detected in significantly fewer cells on average (∼567 vs. ∼940 for negative hits) (**Figure S4D,E**). This was not caused by differences in gRNA representation in the plasmid pool (**Figure S4F**). Thus, we hypothesized that this difference was driven by differences in cell proliferation during reprogramming, as positive hits reprogram astrocytes to a non-proliferating cell type. Indeed, cells that received positive hit gRNAs typically displayed higher expression of neuron marker genes, including MAP2, compared to cells with non-targeting or negative hit gRNAs, and lower expression of genes associated with mitotic DNA replication (**Figure 3G**).

### Single TF perturbations drive reprogramming to multiple neuronal and glial cell states

Stem cells can be differentiated into distinct neuronal subtypes by overexpressing different TFs^35^. However, there has not yet been a high-throughput effort to map the subtype specificity of TFs in astrocyte-to-neuron reprogramming. To fill this gap, we grouped perturbations that converge on similar transcriptomes, then mapped these groupings to neuronal subtypes. We first pseudobulked cells that received each gRNA to reduce data sparsity. Upon unsupervised clustering, pseudobulked perturbations separated into two clusters. One group consisted of perturbations made by gRNAs targeting positive hit TFs and the other consisted of negative hits and non-targeting perturbations (**Figure S5A,B**). We next correlated the transcriptomes resulting from each perturbation (**Figure 4A**). In all cases, different gRNAs targeting the same TF gene displayed concordant effects and were strongly correlated. Similarly, all non-targeting gRNAs grouped together alongside weaker perturbations with relatively few DEGs, forming a large correlated block (**Figure S5C**). Other groups of gRNAs targeting different TFs also converged on similar transcriptomes. For example, gRNAs targeting *NEUROG1*, *NEUROD2*, and *INSM1* produced a highly correlated transcriptome. A separate group was formed by *OLIG2, NR2E1*, and *ZSCAN10*, and another consisted of *ZNF276*, *BCL11A*, *FOXA1*, *TFAP2D*, and *EBF3*. To confirm these results, we also grouped perturbations by a significance-weighted fold change of their DEGs (**Figure S5D**). These distinct methods resulted in highly similar groupings (**Figure S5E**).

**Figure 4:**
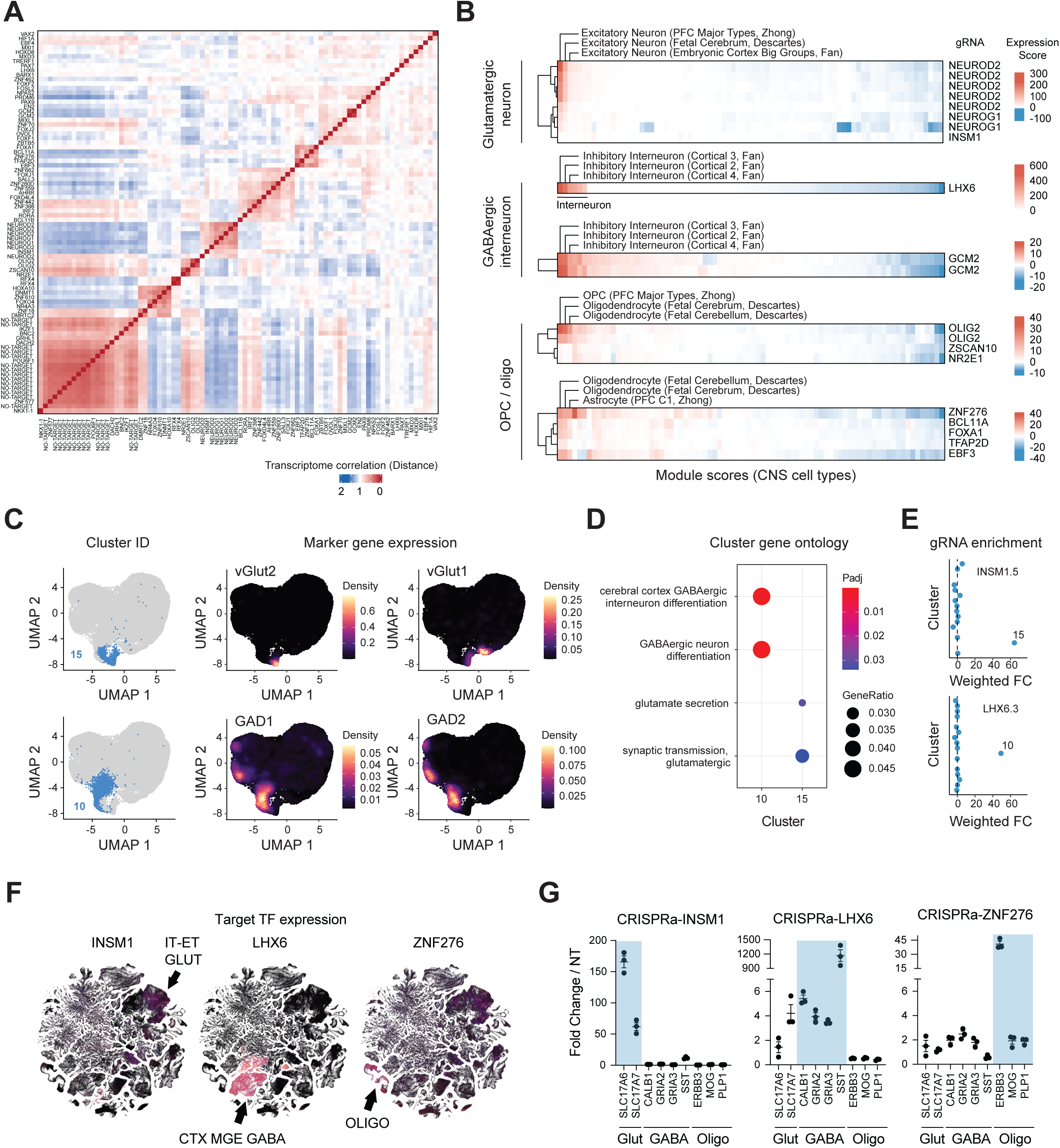
Single TF perturbations promote reprogramming to multiple neuronal and glial cell states. A. Pearson’s correlation-based distance matrix comparing the scaled expression of variable genes. Categories represent transcriptomes pseudobulked by gRNA. All positive hit or non-targeting gRNAs included. B. Module scores of all neural cell type gene signatures from scRNA-seq atlases, ordered from high to low with top 3 scoring modules labeled. Perturbations are grouped on the X axis by similarity in Figure 4A. C. Left: Cells in the relevant UMAP cluster colored in blue. Middle and right: Expression heatmap of key neuronal subtype markers. D. Selected enriched biological processes for neuronal subtype clusters. Statistical significance was determined using a two-tailed Fisher’s exact test followed by Benjamini–Hochberg correction. E. Enrichment for gRNAs in UMAP clusters determined by Seurat’s FindMarkers function with the hurdle model implemented in MAST. Weighted FC = −log10(p_adj_) * abs(fold change). F. Expression of INSM1, LHX6, or ZNF276 in each t-SNE cluster of Allen Brain Cell Atlas^43^. G. Validation of CRISPR perturbations identified to lead to neuronal or oligodendrocyte cell states in Figure 4B. RNA levels of key glutamatergic (glut), GABAergic (GABA), or oligodendrocyte (oligo) marker genes shown. Genes expected to be upregulated by each gRNA highlighted in blue-grey. *p < 0.05, **p < 0.01, ***p < 0.001 by global one-way ANOVA with Dunnett’s post hoc test comparing all groups to non-targeting gRNA.

To determine whether these shared transcriptomes correspond to neuronal subtypes, we re-calculated expression scores for markers of all neural cell types within the Fan, Zhong, or Descartes atlases. As expected, the effects of non-targeting gRNAs showed no enrichment for neural cell subtypes (**Figure S5F**). However, we found that different groupings of targeting gRNAs identified via transcriptome correlation consistently shared top enrichments for specific neural cell types (**Figure 4B**). The group consisting of *NEURODs*, *NEUROG*s, and *INSM1* was enriched for a glutamatergic / excitatory neuronal signature across all three atlases. Other perturbations, notably *LHX6* and *GCM2*, were strongly and consistently enriched for inhibitory interneuron gene signatures. Interestingly, other groups showed enrichment for oligodendrocyte progenitor cell (OPC) and oligodendrocyte gene signatures. OPCs express MAP2 more highly than astrocytes but less than neurons and thus pro-OPC TFs were also captured in the list of hits (**Figure S5G**). Together, these data support reprogramming towards multiple neuronal and glial states after CRISPRa of a single factor.

Based on their strong and specific subtype associations in our dataset, we further examined *INSM1* and *LHX6*. *INSM1* plays key roles in neurogenesis and has been implicated as a downstream node in glia-to-neuron reprogramming^36–38^. *LHX6* has been used to differentiate iPSCs to forebrain GABAergic neurons^39,40^. However, the lineage-specificities of *INSM1* and *LHX6* in astrocyte reprogramming have not previously been defined. In the overall cell distribution, glutamatergic/excitatory neurons were represented by cluster 15, which highly expressed canonical glutamatergic neuron markers vGlut1/2 (encoded by *SLC17A7 / SLC17A6*) and is responsible for transporting glutamate into synaptic vesicles^41^. GABAergic/inhibitory neurons were represented by cluster 10, which highly expressed canonical markers GAD1/2 encoding glutamic acid decarboxylase which is essential for GABA production^42^ (**Figure 4C**). In both cases, cluster marker genes were enriched for terms relating to their respective neuronal subtype (**Figure 4D)**. Consistent with our atlas associations, cells which received a *INSM1* gRNA were strongly and specifically enriched in excitatory neuron cluster 15, while cells which received a *LHX6* gRNA were similarly enriched in inhibitory neuron cluster 10 (**Figure 4E**).

In some but not all cases, these subtype associations were corroborated by in vivo expression patterns established from single cell profiling of human brain tissue (**Figure 4F**). *INSM1* is moderately expressed in multiple cell types in the brain, including but not limited to intratelencephalic and extratelencephalic glutamatergic neurons (IT-ET GLUT)^43^. Furthermore, *INSM1* is also expressed transiently in late-stage neuronal progenitor cells and newborn neurons and therefore would likely be missed if target cell type expression was used as the criteria for reprogramming candidate selection^44^. In contrast, *LHX6* shows clear and specific expression in cortical and pallial neurons from the medial ganglionic eminence (CTX–MGE GABA). Additionally, *ZNF276,* nominated as a target for reprogramming to OPC/oligodendrocyte fate, is expressed most highly in oligodendrocytes and is a driver of oligodendrocyte differentiation in mouse OPCs^45^. Indeed, when activated individually, *INSM1* upregulated glutamatergic markers, *LHX6* upregulated markers of SST+ inhibitory GABAergic interneurons, and *ZNF276* upregulated some but not all tested markers of oligodendrocytes (**Figure 4G**).

### *INSM1* has pro-neuronal activity across systems, cell types, and species contexts

We next tested these factors in other contexts—including different modalities, time points, cell types, and species—to determine if they were context-specific or had robust and broad neurogenic activity.

*Orthogonal modalities:* To test reprogramming with a non-CRISPR-based method, we individually overexpressed an open reading frame (ORF) encoding each annotated splice isoform of INSM1, LHX6, and ZNF276 (**Figure 5A**). As expected, overexpression of the single canonical INSM1 isoform upregulated all tested neuronal markers including post-mitotic neuron marker NeuN. LHX6 has multiple splice isoforms. Neuronal markers DCX and MAP2 were upregulated by all isoforms tested with the exception of isoform 4 (NM_001242335), which lacks the LIM-type zinc finger domain present in the other isoforms. We did not observe upregulation of NeuN after LHX6 ORF overexpression, potentially due to a lack of maturity of induced neurons. ZNF276 upregulated MAP2 expression but did not upregulate neuron markers DCX and NeuN as expected, as these factors are not expressed in oligodendrocytes.

**Figure 5:**
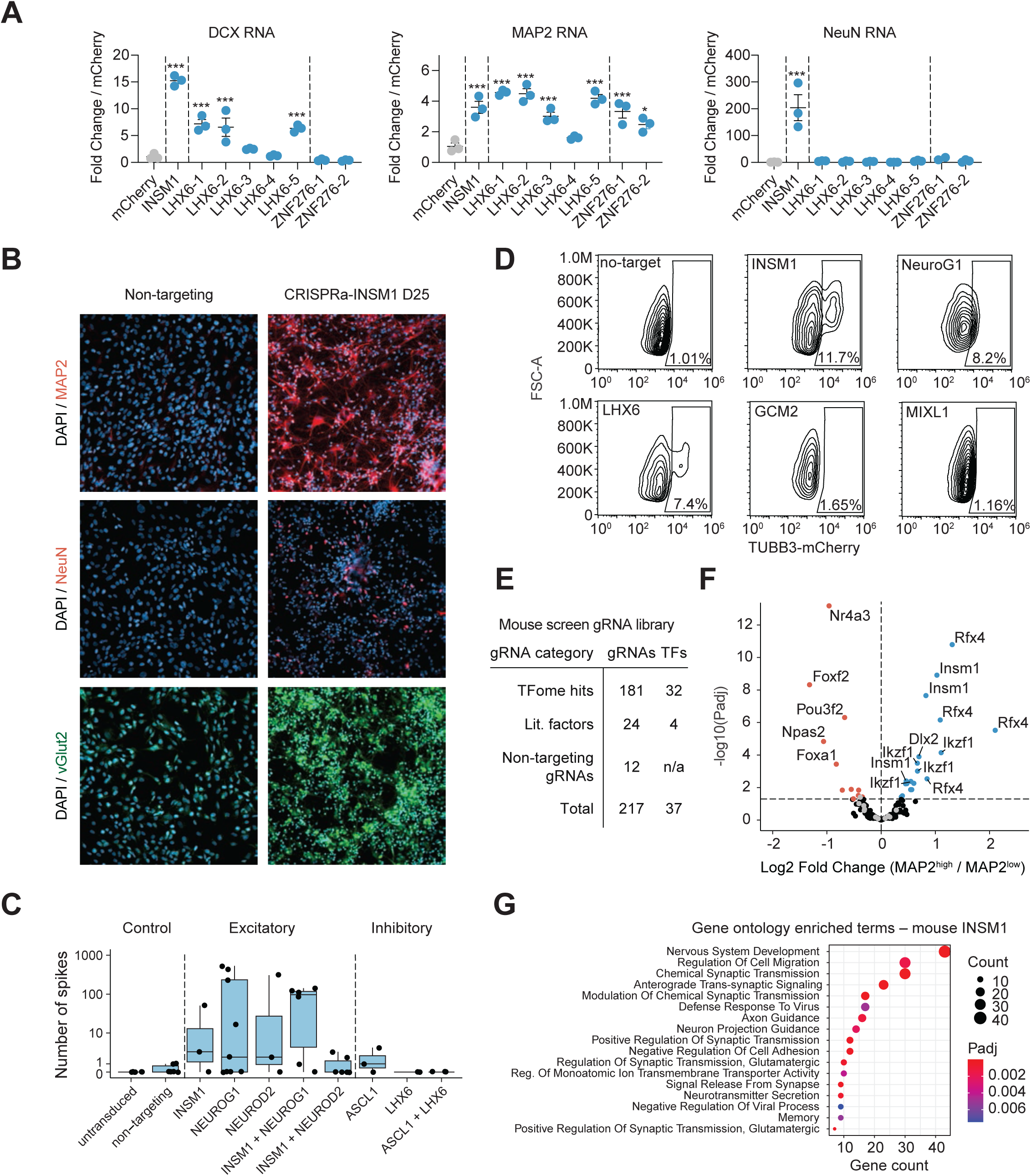
TF candidates generate neuron-like cells from multiple systems and cell types. A. ORF overexpression of annotated INSM1, LHX6, and ZNF276 isoforms. RNA levels of neuron marker genes shown. *p < 0.05, ***p < 0.001 by global one-way ANOVA with Dunnett’s post hoc test comparing all groups to mCherry. B. Immunofluorescence staining of reprogrammed cells to assess MAP2, NeuN, and SLC17A7 expression 25 days after transduction. C. Number of spikes (neuronal firing events) in 5-minute multi-electrode array recording 32 days post-transduction. D. Flow analysis of TUBB3-2A-mCherry expression expression as proxy for early neuronal differentiation 5 days post-transduction of ^VP64^dSpCas9^VP^^64^ iPSCs with gRNA. E. Summary of mouse gRNA sublibrary. F. Significance (P_adj_) versus fold change in gRNA abundance between MAP2-high and MAP2-low populations in mouse CRISPRa screen. G. Top enriched biological processes for upregulated DEGs (L2FC >1, p_adj_ <0.01 determined by DESeq2 vs mCherry, n=121 genes) from RNA-seq 10 days after INSM1 ORF overexpression. Statistical significance of term enrichment was determined using a two-tailed Fisher’s exact test followed by Benjamini–Hochberg correction.

*Extended culture:* Neurons derived from iPSCs display increased cell maturity with prolonged time in culture^46^. Therefore, we sought to test whether astrocytes reprogrammed with CRISPRa-*INSM1* displayed similar dynamics upon extended culture. Immunofluorescent imaging at day 25 post-transduction revealed robust cultures of neuron-like cells positive for markers of post-mitotic glutamatergic neurons (**Figure 5B**). To assess functionality of reprogrammed cells, we used a multielectrode array (MEA) system to monitor electrophysiological activity. Reprogrammed cells displayed functionality after in vitro maturation, firing in a manner consistent with putative excitatory or inhibitory subtype (**Figure 5C, S6A-D**). Higher firing rates were observed after reprogramming with factors linked to excitatory neurons. Interestingly, the highest average spike rate was observed after co-activating *NEUROG1* and *INSM1*, indicating that paired activation increases the proportion of cells that become functional.

*Cell types and species:* Finally, we asked whether factors activated neurogenic differentiation across cell types and species. We activated screen hits in a hiPSC line expressing ^VP64^dSpCas9^VP64^ and tagged with a fluorescent marker inserted into exon 4 of TUBB3, an early pan-neuronal marker gene^47^. This resulted in neuronal differentiation with some but not all TFs, indicating that different TFs and gene regulatory networks drive neuronal cell differentiation from astrocytes compared to pluripotent cells (**Figure 5D**). Both *INSM1* and *LHX6* displayed clear TUBB3+ populations in hiPSCs, suggesting pro-neuronal activity shared between directed differentiation of iPSCs and direct reprogramming of astrocytes. To determine if this neuronal activity was conserved across species, we next screened a library of gRNAs in primary mouse cortical astrocytes (**Figure S6E**). The library consisted of gRNAs targeting the TSS of mouse genes encoding TFs that were hits in the human astrocyte screen and have an ortholog in the mouse genome as well as previously identified additional candidates *Dlx2*, *Nr4a2*, *Atoh1*, and *Lmx1a*^5,10,48,49^ (**Figure 5E, S6F, Supplementary Table 4**). In total, 19 gRNAs were enriched in the MAP2-high bin, largely concentrated across 4 high-confidence hits: *Insm1* (4 of 5 gRNAs), *Ikzf1* (4 of 10 gRNAs), *Rfx4* (4 of 5 gRNAs), and *Dlx2* (3 of 5 gRNAs) (**Figure 5F, S6G, Supplementary Table 4**). Gene ontology analysis of upregulated genes after RNA-seq of mouse astrocytes reprogrammed with *Insm1* revealed broadly neuronal terms, including terms related to nervous system development, chemical and glutamatergic synaptic transmission, axon and neuron projection guidance, and neurotransmitter secretion (**Figure 5G**, **S6H**). Taken together, these data highlight that *INSM1* activation has broad pro-neuronal activity across modalities, cell types, and species, and can produce functional neurons after in vitro maturation.

### Paired gRNA screens identify cofactors of differentiation and subtype specification

CRISPRa-*INSM1* alone reprograms astrocytes to glutamatergic neuron-like cells. However, during development cell fate is specified by coordinated networks of transcription factors^50^. Further, perturbing multiple TFs in combination often leads to improved differentiation and reprogramming^47,51–57^. Therefore, we set out to find cofactors that cooperate with *INSM1* to enhance reprogramming efficiency and glutamatergic subtype specification. We transduced primary human astrocytes with lentivirus encoding ^VP64^dSpCas9^VP64^ and *INSM1* gRNA at high MOI alongside a gRNA library at low MOI and conducted two parallel screens. We screened on levels of 1) MAP2 protein to find factors that enhance reprogramming efficiency, and on 2) SLC17A7 RNA (which encodes vGlut1) using HCR-FlowFISH^58^ to find factors that enhance glutamatergic neuron fate specification (**Figure 6A, S7A**). The screened library consisted of genes encoding hit TFs from the single factor MAP2 screen (**Fig. 2C**), factors from the literature previously implicated in neuronal subtype specification, and controls (**Figure 6B, S7B**). As expected, SLC17A7-targeting gRNAs were the most enriched gRNAs in the SLC17A7 high bin. Additionally, these screens uncovered factors that enhance reprogramming efficiency including IKZF1, *NR4A3*, and *MBNL2*, and factors that increase both efficiency and subtype specificity including *FOXN2*, *TRERF1*, and *NEUROD2* (**Figure 6C, S7C, Supplementary Tables 5,6**). We also performed a screen to discover cofactors that cooperate with *LHX6* to enhance reprogramming efficiency, identifying *FOXJ1* as a top hit (**Figure S7D-H, Supplementary Table 7**).

**Figure 6:**
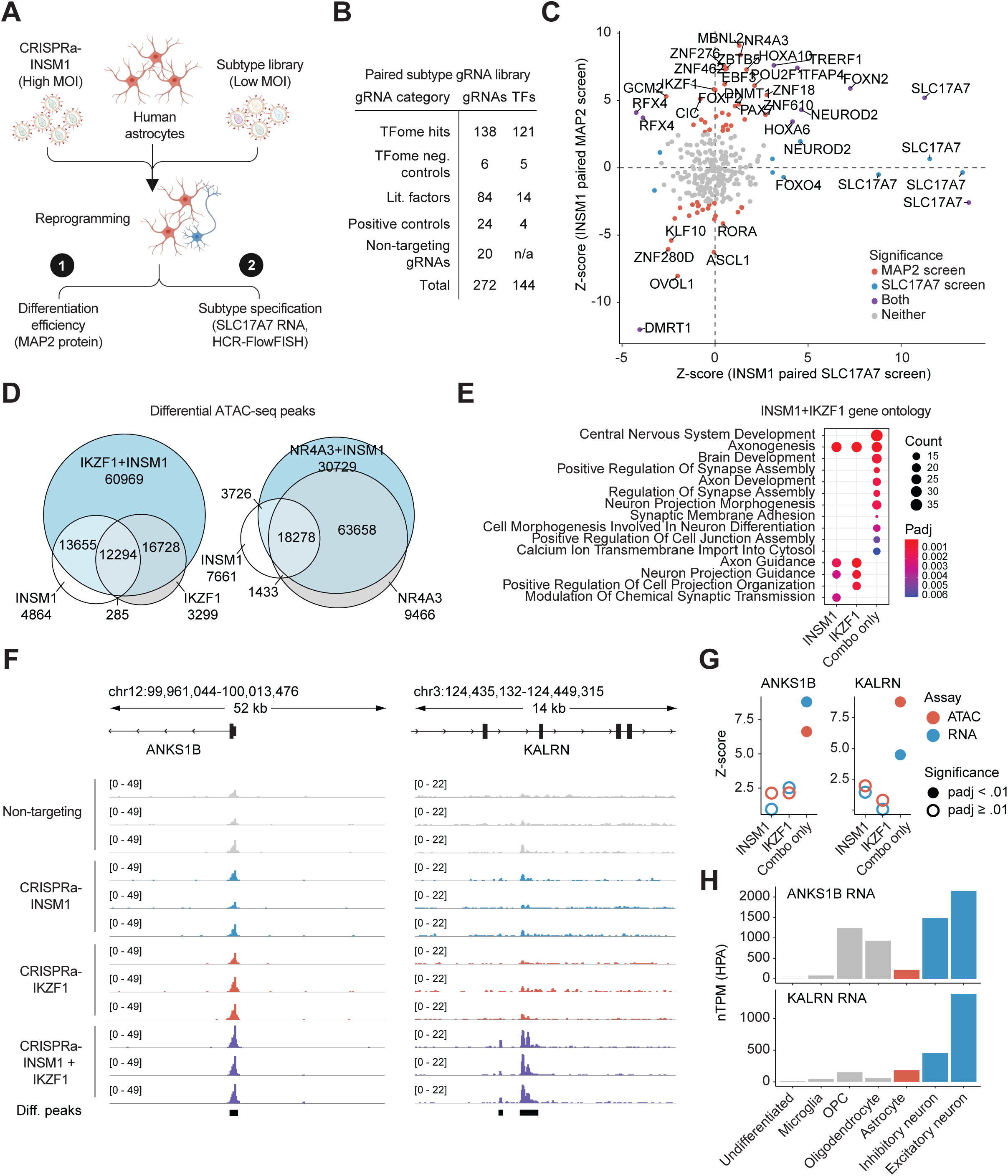
Paired gRNA screens identify cofactors of differentiation and subtype specification. A. Schematic of paired CRISPRa screens. B. Summary of TFpaired screening library. C. Scatter plot of z-score of gRNA abundance in the INSM1 MAP2 paired screen and the INSM1 SLC17A7 paired screen. D. Euler diagrams of differentially accessible peaks. Differential peaks (p_adj_ <.01) for each sample were determined by DESeq2 vs. non-targeting gRNA. E. Top enriched biological processes for genes nearest top 1000 differentially accessible peaks by z-score for INSM1 (I), IKZF1 (IK), or peaks unique in only the combination of INSM1-IKZF1 (I+IK). Statistical significance of term enrichment was determined using a two-tailed Fisher’s exact test followed by Benjamini–Hochberg correction. F. Browser tracks of ATAC-seq (reads per kilobase per million mapped reads [RPKM]-normalized BigWig, bin size = 25bp. ‘Diff.peaks’ denotes peak significance between IKZF1-INSM1 and non-targeting using DESeq2. G. ANKS1B and KALRN peak accessibility and RNA expression indicate lack of significance after reprogramming with individual factors but significance after reprogramming with combination. H. Expression level ofANKS1B and KALRN in neural cell types in the Human Protein Atlas Single Cell Type data^107^.

We next further investigated INSM1 cofactors *IKZF1* and *NR4A3* based on their strong activity in the single factor screens. *IKZF1* was a relatively weak hit as a single factor in human astrocytes (151 DE genes in perturb-seq) but a high-confidence hit in the mouse astrocyte screen, while *NR4A3* was a strong hit in human astrocytes (1914 DE genes) but a negative hit in the mouse astrocyte screen. To further elucidate the interaction between *INSM1* and these cofactors, we conducted ATAC-seq to profile chromatin accessibility of cells reprogrammed with either single factors or TF pairs (**Supplementary Table 8**). As expected, CRISPRa of *INSM1*, *IKZF1*, and *NR4A3* resulted in significant chromatin remodeling at promoters of the targeted genes (**Figure S8A**). Analysis of other differentially accessible peaks revealed that *INSM1* and *IKZF1* have synergistic effects: Alone, they result in 31,098 and 32,606 differential peaks respectively, but when activated together they result in 106,206 differential peaks with 60,969 peaks only differential when both factors are activated (**Figure 6D**, **S8B**). Similarly *INSM1* and *NR4A3* had a synergistic effect when co-activated (120,051 differential peaks) (**Figure S8C**).

Across all tested perturbations, genes nearby differentially accessible peaks were associated with neuron-related biological processes (**Figure S8D**). While genes nearby peaks differential after activation of *INSM1* alone and *IKZF1* alone were associated with axonogenesis, axon guidance, and neuron projection guidance, genes nearby peaks that were only differential when both factors were activated were associated with additional exclusively neuronal gene ontology terms including CNS development, regulation of synapse assembly, neuron projection morphogenesis, and others (**Figure 6E**). Thus, *INSM1* and *IKZF1* alone are pro-neuronal but co-activation of both leads to chromatin remodeling proximal to a large neuronal gene set not significantly impacted by individual perturbations.

### Targeted co-activation of *INSM1* and *IKZF1* remodels chromatin at promoters and intronic regions of key synaptic genes

We next examined specific genes to understand details of chromatin remodeling and associated transcriptional changes after synergistic activation of *INSM1* and *IKZF1*. We focused on genes with important roles in synaptic biology and known disease associations. As a representative example, *ANKS1B* encodes a postsynaptic effector associated with synaptic plasticity and long-term neuronal changes. It is one of the most abundant proteins in the neuronal synapse and has emerged as a top finding in GWA studies examining response to CNS-targeting therapies^59,60^. Further, decreased *ANKS1B* expression is linked to neurodevelopmental syndrome in transgenic mouse models^61,62^. We observed a significant increase (DESeq2 padj = 6.17e-10) in the accessibility of *ANKS1B* promoter after reprogramming with *INSM1* + *IKZF1* despite no significant changes after individual activation of *INSM1* or *IKZF1* (**Figure 6F**). Chromatin changes at key synaptic genes after *INSM1* + *IKZF1* reprogramming were not limited to promoters (**Figure S8E**). For example, we observed significantly increased accessibility of peaks in intronic regions of synaptic gene *KALRN*, a key factor in synaptic plasticity, formation of dendritic projections, and excitatory synaptic transmission. *KALRN* is a central regulator of synaptic function and its dysregulation is linked to a range of neurological disorders^63,64^. Coding variants in *KALRN* have been associated with schizophrenia, highlighting the importance of proper expression levels^65^. These differential peaks were significantly increased only after reprogramming with *INSM1* + *IKZF1 (***Figure 6F**), which similarly led to a matched increase in gene expression levels (**Figure 6G**). Finally, observed increases in expression level were consistent with cell type-specific expression profiles, where *ANKS1B* and *KALRN* are expressed most highly in excitatory neurons compared to other neural cell types (**Figure 6H**). Taken together, these data demonstrate that *INSM1* and *IKZF1* synergistically reprogram human astrocytes and remodel chromatin near key synaptic genes.

### Network analysis of CRISPRa-INSM1 and CRISPRa-IKZF1 reveals mechanisms of interaction and reprogramming

We next sought to identify mechanisms through which *INSM1* and *IKZF1* interact and elucidate the downstream regulatory TFs important for astrocyte-to-neuron reprogramming. After defining differential peaks in response to treatment with either single factor, we separated potential direct targets of *INSM1* or *IKZF1* from indirect targets (targets of downstream TFs) based on whether the differentially accessible regions contained the recognition motif of *INSM1* and/or *IKZF1* DNA binding domains (**Figure 7A**). We found that direct targets of *INSM1* are significantly more abundant in regions opened by CRISPRa-*IKZF1* compared to all peaks (1.53x fold enrichment) (**Figure 7B**). Interestingly, the reverse was not true: Direct targets of *IKZF1* were not enriched in peaks opened by *INSM1*. Further, mapping the genomic locations of these *INSM1* direct targets revealed strong enrichment in promoters (>3x fold increase) compared to all peaks with the *INSM1* recognition motif (46.3% vs 14.0%, **Figure 7C**). Thus, *IKZF1* may enhance *INSM1* activity by preferentially increasing accessibility of its target sites at gene promoters.

**Figure 7:**
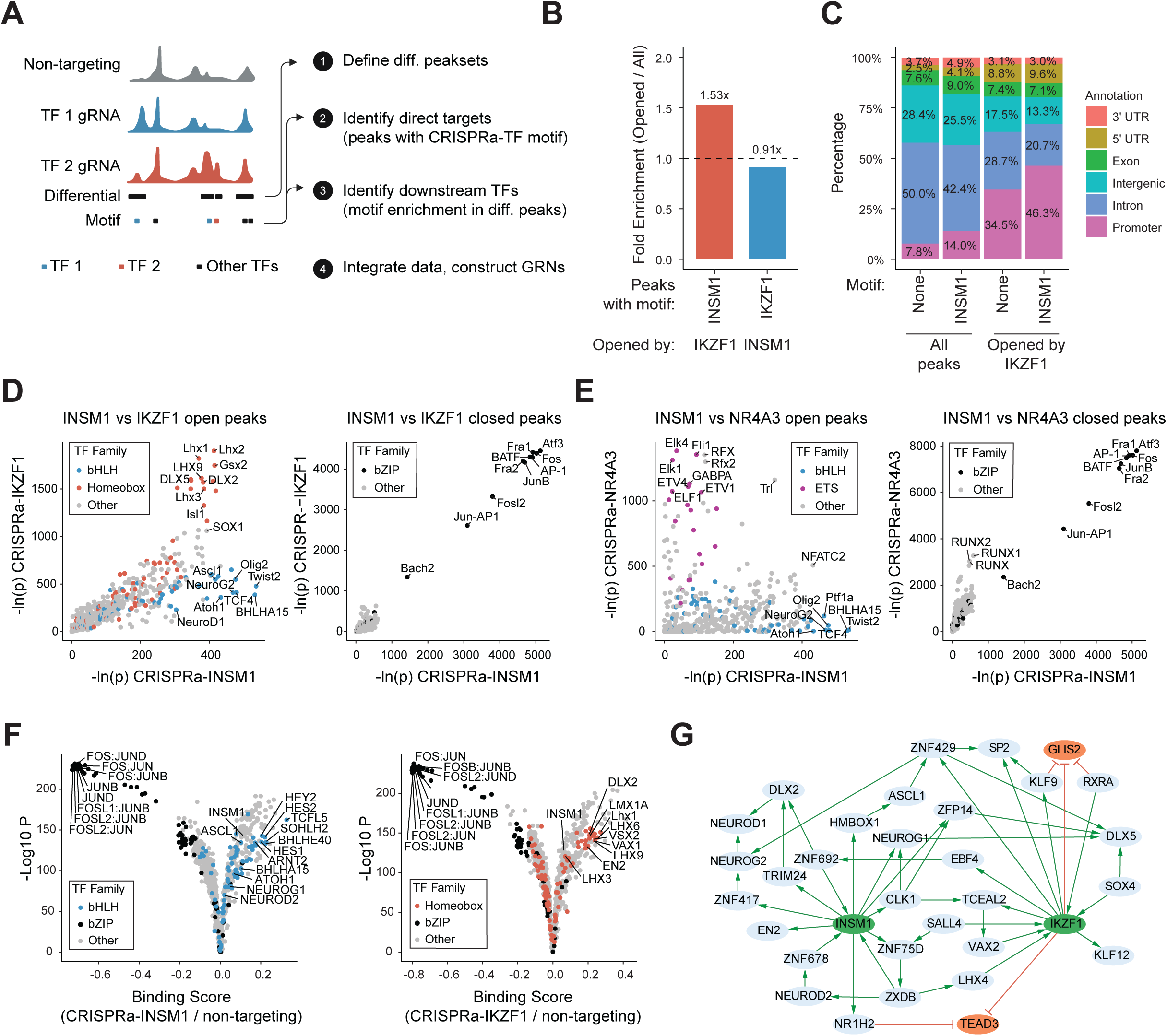
Network analysis of CRISPRa-INSM1 and CRISPRa-IKZF1 reveals mechanisms of interaction and reprogramming. A. Schematic of workflow for identifying direct targets of activated TFs and ‘indirect’ targets of downstream regulators. B. Fold enrichment of prevalence of peaks with INSM1 or IKZF1 motifs across peaksets. Enrichment was calculated as percent of peaks opened by TF1 (but not TF2) with TF2 motif / all peaks with TF2 motif. Set sizes: All peaks: 264,316. With INSM1 motif: 19,854 (7.51%). With IKZF1 motif: 40,076 (15.16%). Peaks opened by IKZF1: 10,539. With INSM1 motif: 1,215 (11.5%). Peaks opened by INSM1: 5,978. With IKZF1 motif: 826 (13.8%). C. Location of peaks across genomic elements. D. Significance of motif enrichment in CRISPRa-INSM1 vs CRISPRa-IKZF1 cells in differentially open regions (left) or differentially closed regions (right). E. Significance of motif enrichment in CRISPRa-INSM1 vs CRISPRa-NR4A3 cells in differentially open regions (left) or differentially closed regions (right). In D and E, HOMER computes P values from the cumulative hypergeometric distribution and does not adjust for multiple hypotheses. Colors indicate transcription factor family. F. Significance versus differential binding of TF motifs (estimated by TOBIAS^66^. Colors indicate transcription factor family. G. A subset of the regulatory connections identified by gene coexpression networks pruned with motif and TF occupancy data (Methods). This subnetwork focuses on TFs that link INSM1 and IKZF1 activity and elucidate downstream neurogenic regulators.

Next, we conducted TF motif enrichment analysis to gain insight into the downstream factors responsible for reprogramming. Overall, the regions with increased accessibility in response to *INSM1* and *IKZF1* displayed correlated motif enrichment. However, bHLH TF family motifs were most strongly enriched in differentially accessible regions after *INSM1* activation while Homeobox TF family motifs were most strongly enriched in differentially accessible regions after *IKZF1* activation (**Figure 7D**, left). In contrast, *INSM1* and *NR4A3* showed orthogonal activity without clear motif enrichment correlation, and motif enrichment in differentially accessible regions after *NR4A3* activation showed the strongest enrichment for ETS TF family motifs (**Figure 7E**, left). Across all perturbations, bZIP TF family motifs including multiple AP-1 family members were strongly enriched in differentially closed regions (**Figure 7D,E**, right). Taken together, these data suggest that *INSM1*, *IKZF1*, and *NR4A3-*based reprogramming function through different downstream regulators, and suggests astrocyte homeostasis may be regulated by a shared mechanism mediated by bZIP family TFs.

Motif enrichment analysis can nominate downstream regulators but does not provide information about whether these motifs are occupied by their respective transcription factors. We therefore used TF footprinting of ATAC-seq data to assess differential TF binding before and after reprogramming^66^. Indeed, cells reprogrammed with CRISPRa-*INSM1* displayed a significant increase in bHLH TF binding to motifs (**Figure 7F**). Similarly, cells reprogrammed with CRISPRa-*IKZF1* displayed a significant increase in binding among some Homeobox TFs, including DLX2, VAX2, EN2, and LHX9. Both populations displayed a significant decrease in bZIP binding, supporting the motif-based results. Notably, CRISPRa-*IKZF1* also increased binding activity of INSM1 to nearly the extent of direct activation of *INSM1*.

Dynamic cellular processes including stem cell differentiation and reprogramming involve finely-tuned complex gene regulatory networks (GRNs). Finally, we integrated these data to map the GRN underlying *INSM1* and *IKZF1*-based astrocyte-to-neuron reprogramming and identify mechanistic relationships (**Figure 7G**). Network analysis suggested a complex interplay between *INSM1* and *IKZF1*, where IKZF1 regulates INSM1 through EBF4 and ZNF692 and both lead to activation of TFs including NEUROG2, SP2 and DLX2, all of which play important roles in neurogenesis^67–69^.

## DISCUSSION

In this study, we established CRISPRa as an approach for human astrocyte to neuron reprogramming. During development, stem cells differentiate into the hundreds of distinct cell types necessary for mammalian organogenesis. This complex cell fate cascade is specified epigenetically through the coordinated actions of transcription factors and their target genomic regulatory elements^70^. We sought to mimic this by using CRISPRa-based reprogramming methods at the endogenous promoter of target genes where activation occurs within the endogenous genomic context with enhancers, introns, UTRs and other regulatory elements intact. Indeed, we found that CRISPRa of bHLH TFs *NEUROD1* and *NEUROG2* led to higher overall expression of downstream neuronal marker genes than cDNA overexpression of the same factors, despite cDNA leading to orders-of-magnitude higher expression of the corresponding TF (**Figure 1E-F**). This highlights the utility of CRISPR-based approaches for engineering complex cellular phenotypes involving complex epigenetic regulation.

We next took advantage of CRISPR’s amenability to high-throughput screening to systematically profile 1,627 genes encoding human TFs for their ability to reprogram primary human astrocytes to neurons. To our knowledge, this is the first high-throughput functional mapping of astrocyte-to-neuron reprogramming factors. This screen uncovered >60 factors able to promote neuronal reprogramming. This list included known neurogenic TFs but also factors not previously tested in this context. Additionally, this work was conducted using human primary cells rather than mouse cells which have been the primary focus in this area. Mouse cells are more readily reprogrammed, resulting in higher efficiencies than their human counterparts^53,71^. Thus, human cells may require additional or alternate factors to achieve similar reprogramming efficiencies due to the broad differences between murine and human neurobiology and plasticity.

The large number of hits in this screen highlights the potential cellular plasticity of astrocytes. In vivo, astrocytes are responsive to their microenvironment and can undergo drastic molecular and phenotypic changes including proliferation and scar formation after injury or disease. This process, referred to as reactive astrocytosis, is an area of active investigation and is increasingly appreciated to be context-dependent and heterogeneous^12,72–76^. While further work is needed to elucidate its impact on neurodegeneration in different contexts, proliferative reactive astrocytes represent an attractive starting material for neuronal reprogramming. To accomplish this, it will be important to further develop delivery methods to specifically target reactive astrocytes in vivo^77^.

Profiling of hits that emerged from the TFome CRISPRa screen by scRNA-seq revealed that activation of some TFs resulted in highly correlated transcriptomes, which in turn were enriched for neuronal subtype-specific gene signatures from pulic scRNA-seq brain tissue atlases. Among others, these results nominated *INSM1* as inducing pro-excitatory glutamatergic neurons and *LHX6* and *GCM2* as promoting pro-inhibitory GABAergic interneurons. Our TFome flow cytometry-based screen was conducted on MAP2 expression due to its established role as a pan-neuronal marker gene. MAP2 is also expressed to a lower level in OPCs^30^ (**Figure S5G**). We also uncovered perturbations enriched for OPC/oligodendrocyte gene signatures, including 2 gRNAs targeting *OLIG2*, a master regulator of oligodendrocyte fate^78^ and *ZNF276,* which has been linked to OPC differentiation^45^ but has not been previously tested in astrocytes.

To determine whether reprogramming factors we identified were specific to human astrocytes or had broad pro-neuronal activity, we tested them in iPSCs and primary mouse cortical astrocytes. *INSM1* in particular displayed broad neurogenic activity in these contexts, differentiating a subpopulation of human iPSCs to TUBB3-positive cells and resulting in multiple *Insm1*-targeted gRNAs enriched in the MAP2-high bin in a primary mouse astrocyte reprogramming screen. *Rfx4*, *Ikzf1*, and *Dlx2* also emerged as high-confidence hits in this mouse screen, consistent with previous reports demonstrating their robust neurogenic activity^10,79,80^. However, other TFs such as *GCM2* did not display pro-neuronal activity in other cell types, potentially indicating cell type- or species-specific GRNs constraining reprogramming.

During development, cell fate is specified through coordinated interacting networks of TFs. Hence, we sought to discover cofactors that cooperate with *INSM1* to enhance neuronal differentiation and glutamatergic subtype specification. We further investigated TF combinations *INSM1*-*IKZF1* and *INSM1*-*NR4A3* due to the activity of *IKZF1* and *NR4A3* in our other screens. ATAC-seq of astrocytes reprogrammed with single TFs or combinations revealed that *IKZF1* and *NR4A3* synergize with *INSM1* in different ways. The effect of *INSM1*-*IKZF1* is greater than the sum of individual perturbations (106,206 DA peaks vs 63,704 DA peaks, **Figure S8B**). The unique effect of *INSM1*-*IKZF1* combination is pro-neuronal, indicated by a neuronal gene module not remodeled by either individual TF (**Figure 6E**) including peaks at important synaptic promoters and intronic regions (**Figure 6G**). In contrast, the number of differentially accessible peaks after reprogramming with *INSM1*-*NR4A3* was more aligned with the sum of peaks from individual perturbations (**Figure S8B**). Further, we elucidate modes of TF interaction and downstream regulators leading to astrocyte-to-neuron reprogramming (**Figure 7**). The mechanism of one TF potentiating the effect of a second TF by increasing accessibility of a subset of potential binding sites, as we have observed with IKZF1 and INSM1, has been observed in other contexts including pluripotent reprogramming^81^ and is likely a general mode of lineage specification that may be more broadly repurposed for reprogramming applications.

Overall, through this study we have identified an expanded list of TFs to be targeted for astrocyte-to-neuron reprogramming, broadly expanding the toolbox for cell reprogramming-based neuroregenerative therapies. Our screens conducted in primary neural cells were enabled by the use of all-in-one lentiviral vectors with robust activation activity and intracellular antibody and FISH-based FACS screen readouts. These features eliminated the need for the generation and expansion of cell lines with stable dCas9 expression or a reporter gene, which would not have been feasible with primary human astrocytes. We expect that this approach can be applied more widely to study relevant phenotypes including reprogramming in other primary cell types with limited proliferative potential in vitro.

## METHODS

### Animal use

All animal experiments were conducted with strict adherence to the guidelines for the care and use of laboratory animals of the National Institutes of Health. All mice and rats were used in accordance with the Institutional Animal Care and Use Committee (IACUC) and the Duke Division of Laboratory Animal Resources (DLAR) oversight (IACUC Protocol Number A217-21-10). All mice were housed under typical day/night conditions of 12-hour cycles.

### Plasmids

Plasmids were constructed using Gibson assembly (NEB). The all-in-one CRISPRa lentiviral plasmid expressing ^VP64^dSpCas9^VP64^, a gRNA scaffold, and a puromycin selection cassette used most widely in this work was generated by modifying Addgene plasmid #71236 to replace dSpCas9-KRAB domain with ^VP64^dSpCas9^VP64^. This plasmid was further modified to generate an all-in-one CRISPRa lentiviral plasmid expressing ^VP64^dSpCas9^VP64^, a gRNA scaffold, and Thy1.1 by replacing PuroR with Thy1.1. Individual gRNAs were ordered as oligonucleotides (IDT), phosphorylated, hybridized, and cloned into the plasmids using BsmBI sites. cDNA overexpression plasmids were ordered from Addgene (ASCL1: #162345; NeuroD1 #162338) or cloned. NeuroG2 cDNA-overexpression plasmid was generated by modifying Addgene plasmid #162345 to replace ASCL1 cDNA with NeuroG2 cDNA from a gBlock (IDT). INSM1, LHX6 and ZNF276 overexpression plasmids were constructed by amplifying ORFs from the Multiplexed Overexpression of Regulatory Factors (MORF) Library^82^ (Addgene #192821) using ORF-specific primers or ordered from Addgene (INSM1 #144140).

### Human astrocyte culture

Primary human astrocytes isolated from the cerebral cortex (Sciencell 1800) were purchased and cultured according to the manufacturer’s instructions. Briefly, cells were grown on poly-d-lysine (PDL)-coated plates in Astrocyte Medium (Sciencell 1801) supplemented to 10% FBS (Gibco). Media was refreshed every 2 days until cultures were 70% confluent, then every 3 days until cultures reached 95% confluence, at which point they were passaged with 0.025% Trypsin. Astrocytes were grown for two passages to differentiate potential contaminating progenitor cells and replated to remove potential neurons. All experiments were initiated upon replating after the second passage reached confluence.

### Human astrocyte reprogramming

For the direct reprogramming of human astrocytes to neurons, primary human astrocytes were maintained as described above until the second passage reached confluence. Cells were then dissociated, counted, and seeded at a density of 9 x 10^4^ cells / cm^2^ in media containing packaged lentivirus. Transduction was noted as day 0. Media was refreshed after 24 hours (day 1). On day 2, astrocyte media was supplemented with 1ug/mL puromycin (Thermo Fisher), which was added to all media until day 6. On day 3, astrocyte media was replaced with basal co-culture media consisting of DMEM/F12 (Gibco) supplemented with 0.5% FBS (Gibco), 1x N-2 (Gibco), 3.5mM Glucose (Gibco), and 100 U/mL Penicillin-Streptomycin (Gibco). On day 8, co-culture media was supplemented with 20 ng/ml BDNF and 10 ng/ml NT3 (Peprotech). Media was refreshed every other day.

### Glutamate uptake assay

Supernatant was collected from wells with and without astrocytes and glutamate levels were measured using a fluorometric assay according to the manufacturer’s protocol (Cell Biolabs STA-674). To reverse glutamate transporter activity, cells were treated with 1mM 4-Aminopyridine (4-AP, Millipore Sigma) for 30 minutes. Media was collected 1 day after 4-AP treatment and measured on a Promega GloMax plate reader.

### Lentivirus packaging

Lentivirus was packaged as described in McCutcheon et al. Nature Genetics 2024^83^. Briefly, HEK293T cells were counted and plated in OptiMEM Reduced Serum Medium (Gibco) supplemented with 1x Glutamax (Gibco), 5% FBS (Gibco), 1 mM Sodium Pyruvate (Gibco), and 1x MEM Non-Essential Amino Acids (Gibco). After 12-18 hours, HEK293T cells were transfected with pMD2.G, psPAX2, and transgene using Lipofectamine 3000 (Thermo Fisher). Media was exchanged 6 hours after transfection and lentiviral supernatant was collected and pooled at 24 hours and 48 hours after transfection. Collected supernatant was centrifuged or filtered to remove cellular debris and concentrated to 50x using Lenti-X Concentrator (Takara).

### Lentiviral titration

Titration of packaged lentivirus was carried out according to the protocol outlined in Gordon et al. Nature Protocols 2020^84^. Briefly, cells were transduced with serial dilutions of lentivirus. After 24 hours, media was changed. After four days, cells were rinsed three times with PBS and genomic DNA was extracted with a DNeasy Blood & Tissue Kit (Qiagen). Integrated titer was then determined via qPCR by utilizing primer sets specific to genomic DNA (LP34), integrated viral DNA (WPRE), and unintegrated plasmid backbone (BB). Viral volumes which led to cell death were excluded from analysis.

### RT-qPCR

Total RNA was isolated from transduced primary astrocytes at day 10 with a Total RNA Purification Plus Kit or Total RNA Purification Plus Micro Kit (Norgen) and reverse transcribed using Supercript VILO (Thermo Fisher) with an equal mass input. qPCR was performed with Perfecta SYBR Green Fastmix (Quanta BioSciences). Amplification and measurement were completed using a CFX96 Real-Time PCR Detection System (Bio-Rad). Standard curves were constructed before primers were used for quantification, and amplicon product specificity was confirmed by melt curve analysis.

### Immunocytochemistry

For imaging experiments, the reprogramming protocol described above was conducted on a PDL-coated 8-well µ-Slide (Ibidi Bioscience). Cells were rinsed with PBS and fixed with 4% formaldehyde (Pierce) for 15 minutes at room temperature. Cells were rinsed 3x with PBS and permeabilized with PBS with 0.1% Triton-X (Sigma), rinsed, and blocked with PBS, 0.1% Tween-20 (Sigma), and 10% Normal Goat Serum (Sigma). The following primary antibodies were used with incubations overnight at 4C: Rabbit anti-GFAP (1:500 dilution, Proteintech 16825-1-AP), rabbit anti-MAP2 (1:500 dilution, Abcam ab183830), mouse anti-NeuN (1:1000 dilution, Millipore Sigma MAB377), rabbit anti-NeuN (1:300 dilution, CST 24307) and/or guinea-pig anti-vGlut2 (1:2000 dilution, Millipore Sigma AB2251-I). Cells were rinsed 3x and incubated with DAPI (Invitrogen) and cross-adsorbed secondary antibodies (Invitrogen) conjugated to Alexa Fluor 488 or 647. Cells were rinsed 3x with PBS and imaged with a Zeiss 780 upright fluorescent microscope.

### RNA-sequencing

Total RNA was isolated from transduced primary astrocytes at day 10 with a Total RNA Purification Plus Kit or Total RNA Purification Plus Micro Kit (Norgen). RNA was submitted to Azenta for standard RNA-seq with polyA selection with ERCC spike-in. Libraries were sequenced on an Illumina sequencer (150 cycles, PE). Reads were trimmed using Trimmomatic v0.32^85^ and aligned to GRCh38 human genome using STAR v2.4.1a^86^. Gene counts were obtained via featureCounts (Subread v1.4.6-p4)^87^ using the comprehensive gene annotation in Gencode v22. DESeq2^88^ was used for differential expression (DE) analysis. DESeq2 employs a negative binomial generalized linear model (GLM). DE genes are determined using a Wald test (padj <0.01). Upregulated and downregulated DEGs (|l2fc| > 1) were input into EnrichR’s^89^ GO Biological Processes 2023 database for functional annotation.

### Intracellular flow cytometry

Cells were rinsed with PBS, collected with 0.025% Trypsin, singularized, and resuspended in Intracellular Fixation Buffer (eBioscience) for 20 minutes at room temperature on a rocker. Cells were then rinsed and permeabilized with Intracellular Permeabilization Buffer (eBioscience). Following permeabilization, cells were rinsed and blocked for 10 minutes at room temperature by resuspension in permeabilization buffer with 0.2M glycine (Sigma) and 2.5% FBS (Gibco). Staining: Blocked cells were incubated with rabbit anti-MAP2 antibody conjugated to Alexa Fluor 488 (Abcam ab225316) for 30 minutes at room temperature. Cells were rinsed and sorted and/or analyzed on a Sony SH800z Cell Sorter.

### Construction of gRNA library targeting all human transcription factors

gRNAs targeting each TSS of all putative human TFs^31^ and 487 non-targeting gRNA controls (5% of targeting gRNAs) were subset from an optimized library ^90^. For each TF that had <6 gRNAs present in the referenced library, additional unique gRNAs were subset from an earlier published gRNA library^91^, for a total library size of 10,233 gRNAs. gRNAs were then ordered as an oligo pool from Twist Biosciences and cloned into an AIO CRISPRa vector by Gibson assembly. Adequate representation of all gRNAs was confirmed by Illumina sequencing.

### CRISPRa screening of all human transcription factors

Cell handling and sorting: Primary human astrocytes (n=3 donors) were transduced with lentivirus encoding for ^VP64^dSpCas9^VP64^ and gRNA library at MOI = 0.3 and proceeded through the reprogramming protocol described above. Cells were collected, fixed, permeabilized, and stained for intracellular MAP2 as described above. The lower and upper 10% of MAP2 expression were sorted for subsequent gRNA library construction and sequencing. All replicates were sorted at a minimum of 150x coverage. Genomic DNA isolation, gRNA cassette amplification, and sequencing: Crosslinking was reversed at 65C overnight using a PicoPure DNA Extraction Kit (Arcturus) and DNA was purified by ethanol precipitation. Integrated gRNA cassettes from each sample were then amplified from genomic DNA with barcoded custom i5 and i7 primers for Illumina sequencing (**Supplementary Table 7**). After double-sided SPRI bead selection, barcoded amplicons were pooled, diluted, and sequenced on an Illumina MiSeq using custom Read 1 and index primers (**Supplementary Table 7**). Screen analysis: FASTQ files were aligned to gRNA libraries using Bowtie2^92^. Counts for each gRNA were extracted and used for further analysis. gRNAs with outlier low counts were removed from further analysis. Individual gRNA enrichment was determined using DESeq2^88^ to compare gRNA abundance between high and low expression conditions for each screen. gRNAs with p <.01 were designated as hits.

### Single-cell CRISPR activation screening of TFs of interest

Cell handling and sorting: Primary human astrocytes (n=2 donors) were transduced with lentivirus encoding for ^VP64^dSpCas9^VP64^ and gRNA library consisting of all hits from the FACS-based screen and 14 non-targeting gRNAs at MOI = 0.3 and proceeded through the reprogramming protocol described above. Cells were collected and gRNA and gene expression libraries were prepared using the 10X High-throughput kit with 5’ gRNA Direct Capture (10x Genomics) according to manufacturer protocol and sequenced on an Illumina Novaseq. Demultiplexing and UMI count generation for each transcript and gRNA per cell barcode was performed using CellRanger v6.0.1 (10x Genomics). UMI counts tables were extracted and used for subsequent analyses in R using Seurat v4.1.0^93^ and normalized with sctransform^94^. Low quality cells were discarded. Remaining high-quality cells across donors were aggregated for further analyses. gRNAs were assigned to cells if they exceeded the count threshold recommended by the Cellranger mixture model. Cells were then grouped for differential expression analysis using MAST^95^ based on gRNA identity. DE testing: For differential gene expression analysis, for each gRNA, cells that received a given gRNA were compared to cells that only received a non-targeting gRNA using Seurat’s FindMarkers function with the hurdle model implemented in MAST. Upregulated DEGs were input into EnrichR’s GO Biological Process 2023 database for functional annotation as described above. Module scoring: Module scores for each cell type in published atlases were calculated using MSigDB (Dolgalev. *R package version 7.5.1.9001.* 2022) and applied to cells, pseudocells, or DE gene lists with the AddModuleScore function in Seurat. Pseudobulking and transcriptome correlation: To calculate the overall effect of each perturbation on cell transcriptomes, for each gRNA, the transcriptomes of each cell that only received that gRNA were averaged to create one ‘pseudocell’ per gRNA in the library. Positive hits and non-targeting gRNAs were considered in subsequent analysis. Variable features were scaled and used for PCA or measured with a Pearson’s correlation for transcriptome comparisons.

### Electrophysiology

To measure the electrical activity of reprogrammed cells, astrocytes were replated onto PDL-coated 96-well MEA plates (Axion Biosystems) 4 days after lentiviral transduction. Cells were plated at a density of 100,000 cells per well in a concentrated droplet on the electrode surface to facilitate network formation in the area of measurement. Cells were maintained in co-culture media with BDNF and NT3, which was changed twice per week, including the day before recording. Electrophysiological records were conducted on a Maestro MEA system (Axion Biosystems). Recordings were taken for a 5-minute period after cell viability measurement. Recording data were analyzed using the Axion Neural Metric Tool.

### iPSC culture and neuronal differentiation

iPSCs were cultured and differentiated as previously described^47^. Briefly, TUBB3-2A-mCherry RVR iPSCs expressing ^VP64^dSpCas9^VP64^ were grown on Matrigel (Corning) in complete mTesR (Stemcell Tech). For neuronal differentiation, iPSCs were transduced with lentivirus encoding gRNA cloned into Addgene plasmid #162335 and culture medium was changed to DMEM/F-12 Nutrient Mix (Gibco), 1x B-27 supplement (Gibco), 1x N-2 supplement (Gibco), 1x CultureOne™ Supplement (Gibco), 25 mg/mL gentamicin (Sigma), and 1ug/mL puromycin. Five days after transduction, cultures were analyzed for mCherry expression using a Sony SH800z Cell Sorter.

### Mouse primary astrocyte isolation and culture

C57BL/6 P2 mouse cortices were microdissected and digested in DPBS (Thermo Fisher), 10 units/ml papain (Worthington Biochemicals), and 12,500 units/mL DNaseI (Worthington Biochemicals). Following digestion, tissue was triturated in low-ovomucoid solution consisting of DPBS, 1.5 mg/ml BSA (Sigma), and 1.5mg/ml Trypsin inhibitor (Worthington Biochemicals) with DNase followed by high-ovomucoid solution consisting of DPBS, 4.28 mg/ml BSA (Sigma), and 4.28 mg/ml Trypsin inhibitor (Worthington Biochemicals) with DNase. Astrocytes were then filtered and plated on PDL-coated flasks in mouse astrocyte growth media (mAGM) consisting of DMEM (Gibco 11960), 10% FBS (Gibco), 100 U/mL Penicillin-Streptomycin (Gibco), 2 mM L-Glutamine (Gibco), 5 µg/ml Insulin (Sigma), 1 mM Sodium Pyruvate (Gibco), 5 µg/ml N-Acetyl-L-cysteine (Sigma), and 10uM hydrocortisone (Sigma). Three days after plating, mAGM was replaced with dPBS. Culture flasks were then shaken vigorously by hand until only astrocytes remained attached. Media was then replaced with fresh mAGM. Two days later, 10uM cytosine arabinoside (AraC, Sigma) was added to culture media to remove quickly dividing cells. Two days later (7 days after dissection), cells were replated and transduced for reprogramming.

### Mouse astrocyte reprogramming

Purified mouse cortical astrocytes were replated at a density of 9 x 10^4^ cells / cm^2^ in mAGM containing packaged lentivirus for CRISPR screening or individual TF validation. Media was refreshed after 24 hours (day 1). On day 2, mAGM was supplemented with 1.5ug/ml puromycin (Thermo Fisher), which was added to all media until day 6. On day 3, astrocyte media was replaced with co-culture media consisting of Neurobasal (Gibco) supplemented with 1x B-27 (Gibco), 1x Glutamax (Gibco), and 100 U/mL Penicillin-Streptomycin (Gibco). On day 6, co-culture media was supplemented with 20 ng/ml BDNF and 10 ng/ml NT3 (Peprotech). Media was refreshed every other day until day 14, when cells were harvested for flow cytometry or RNA isolation.

### Paired screening for INSM1 and LHX6 cofactors

Astrocytes were transduced with a lentiviral vector encoding ^VP64^dSpCas9^VP64^-2A-Thy1.1 and INSM1 or LHX6 gRNA at high MOI, and a lentiviral pool encoding the screening library at low MOI. MAP2 screens: After reprogramming, cells were fixed, permeabilized, stained for intracellular MAP2 expression, sorted, and analyzed according to the method described above. SLC17A7 screen: To screen for factors that cooperate with INSM1 to enhance glutamatergic subtype specification, cells were sorted based on abundance of SLC17A7 RNA using HCR-FlowFISH according to the method described in Reilly et al. 2021^58^ with the following modifications: SLC17A7 probeset and buffers were ordered from Molecular Instruments (https://www.molecularinstruments.com), and SLC17A7 probeset was used at a final concentration of 8nM overnight. Briefly, reprogrammed cells were fixed for 10 minutes in PBS, 4% paraformaldehyde (Pierce), 0.1% Tween-20 (Sigma). After washing, cells were resuspended in hybridization buffer, and incubated with 8nM SLC17A7 probe overnight at 37C. The next day, cells were washed in probe wash buffer (Molecular Instruments) then 5x SSCT (sodium chloride sodium citrate with 0.1% Tween). Cells were then incubated with 60nM snap-cooled hairpin in amplification buffer. Cells were incubated overnight at room temperature, washed, then sorted using a Sony SH800z Cell Sorter. Downstream screen processing and analysis was performed as outlined above.

### ATAC-seq

A total of 5×10^4^ nuclei were prepared for Omni ATAC-seq as previously described^96^. Libraries were sequenced on an Illumina NextSeq 2000 with paired-end 50-bp reads. Reads were QCed, trimmed, aligned, and filtered using the ENCODE ATAC-seq pipeline (https://github.com/ENCODE-DCC/atac-seq-pipeline). Peaks were called with MACS2^97^ and optimal narrowPeak files were merged across samples with bedtools merge^98^. A counts table containing the number of reads in peaks for each sample was generated using featureCounts (Subread v1.4.6-p4)^87^ and used for differential analysis with DESeq2^88^. Genes nearest to each peak were assigned using bedtools closest and annotated with biomaRt^99^. Peaks overlapping two or more genes were excluded from analysis. For visualization of peaks, BAM files were normalized for visualization using deepTools bamCoverage^100^. For each sample, differentially accessible peaks were annotated by genomic element type using ChIPseeker R package (v1.38.0)^101^. Genomic annotation categories were prioritized hierarchically as follows: Promoter > UTR > Exon > Intron > Intergenic.

### Motif and transcription factor occupancy analysis

To identify transcription factor (TF) motifs enriched in chromatin regions differentially accessible upon CRISPRa activation of candidate TFs, we performed motif enrichment analysis using HOMER (v4.11)^102^. Differential peaks identified by ATAC-seq analysis were grouped based on directionality of fold-change and specific TF perturbations. Each group of peaks was analyzed using findMotifsGenome.pl with the -size given option to retain exact peak boundaries. Motif enrichment was calculated relative to a unified background set consisting of the union of all accessible peaks from the experiment. Significantly enriched known motifs were reported and interpreted in the context of TF activity, synergy, and putative regulatory network engagement. A TOBIAS-based footprinting analysis was performed to access differential transcription factor (DTF) binding in CRISPRa TF samples and controls. Replicates for each condition were merged using samtools (v1.21)^103^. The resulting BAM files were then sorted and indexed. To correct for Tn5 insertion bias, ATAC-seq BAM files and peak sets were processed by TOBIAS version 0.17.1 ATACorrect with the hg38 reference genome, which generated corrected BigWig files for downstream analysis. TOBIAS FootprintScores was then applied to the corrected files and corresponding union peaks to generate footprint scores for each condition. DTF binding analysis was performed using TOBIAS BINDetect, with motif definitions from the JASPAR 2024 CORE vertebrates non-redundant database in MEME format.

### Gene regulatory network inference

Gene regulatory networks (GRNs) were inferred from bulk RNA-sequencing data generated under IKZF1, INSM1, and IKZF1-INSM1 perturbation conditions using GRNBoost2^104^ via the pySCENIC v0.12.1 pipeline^105^. GRNs were pruned to retain nodes that met at least one of the following criteria: a) differential expression between the perturbation and non-targeting control conditions at adjusted p < 0.01, or b) differential predicted binding between perturbation and non-targeting control conditions at adjusted p < 0.01, as identified by TOBIAS BINDetect^66^. Edges were retained only if both source and target nodes were present in the filtered node set. We visualized the resulting GRNs in Cytoscape^106^ and annotated edges as either ‘activation’ or ‘inhibition based on log2 fold-change in expression of the target gene between conditions (i.e. log2FC ≥ 0 for activation, log2FC < 0 for inhibition). Individual networks for each perturbation condition were further manually curated to reduce visual complexity and highlight regulatory relationships of interest.

### Data availability

All sequencing datasets can be accessed via the NIH Impact of Genetic Variation on Function Data portal at https://data.igvf.org/. Accession numbers are available in the **Supplementary Table**.

## ACKNOWLEDGEMENTS

This work was supported by NIH Grants UM1HG012053, RM1HG011123, T32GM008555, and R01MH125236.

We thank all members of the Gersbach laboratory, Justin Savage, Lucio Schiapparelli, Ruhi Rai, and Alejandro Barrera for technical assistance and helpful discussions. Illustrative schematics were created using biorender.com.

