## Supplementary Figures for "Comprehensive profiling of transcription factors for reprogramming human astrocytes to neuronal cells through endogenous CRISPR-based gene activation"

Figure S1: Reprogramming primary human astrocytes with bHLH transcription factor CRISPRa. Related to Figure 1.

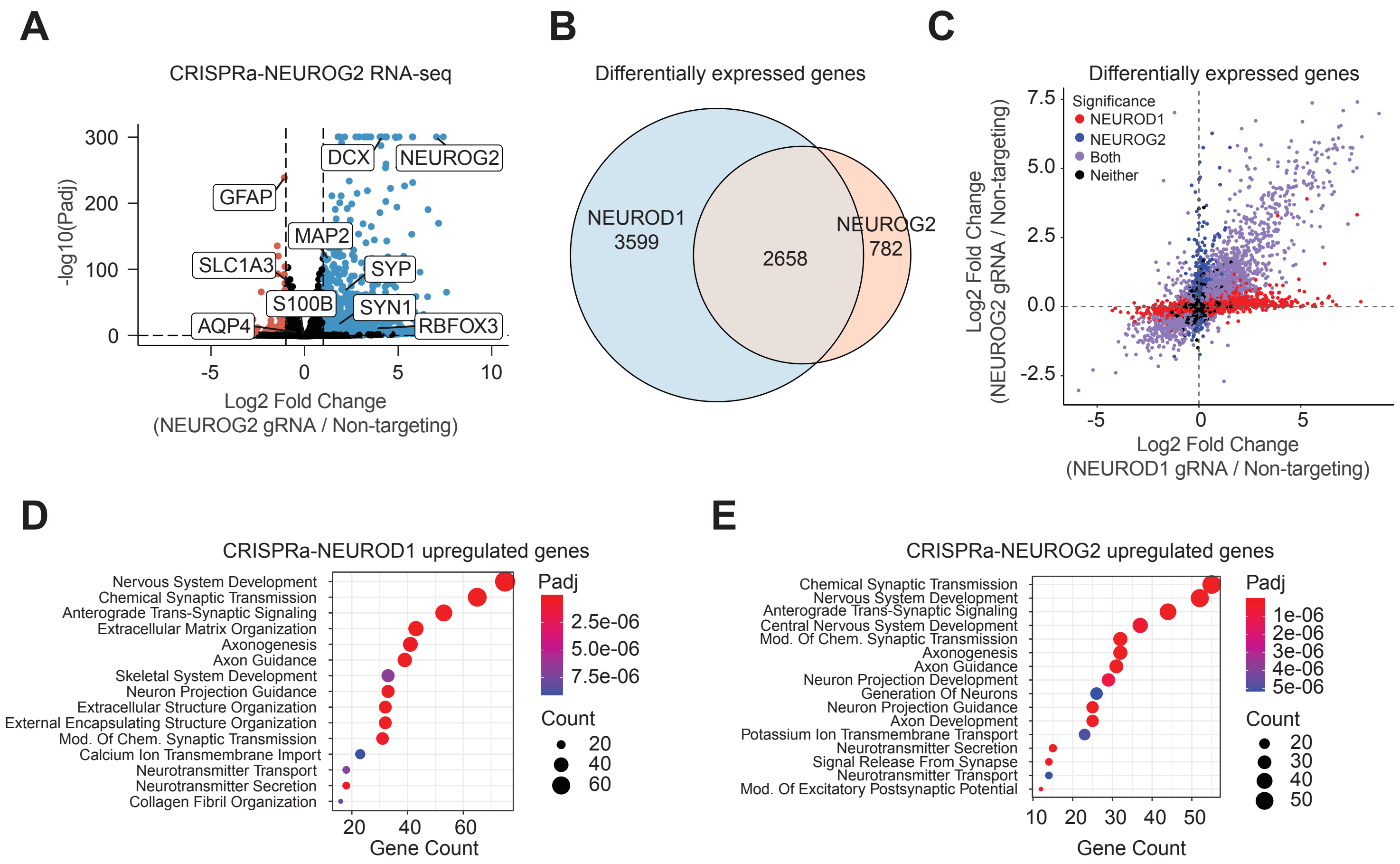

**Figure S1. Reprogramming primary human astrocytes with bHLH transcription factor CRISPRa. Related to Figure 1:**

A. Differential expression analysis of RNA sequencing of $^{VP64}$dSpCas9$^{VP64}$ + NEUROG2 gRNA (CRISPRa-NEUROG2) vs. $^{VP64}$dSpCas9$^{VP64}$ + non-targeting gRNA, 10 days post-transduction. Canonical astrocyte and neuron marker genes are labeled. DE genes are determined using a Wald test by DESeq2 (padj $<0.01$).
B. Overlap of upregulated DE genes after CRISPRa-NEUROD1 and CRISPRa-NEUROG2.
C. Scatter plot comparing fold changes of DE genes after CRISPRa-NEUROD1 and CRISPRa-NEUROG2.
D. Top enriched biological processes for upregulated DE genes after CRISPRa-NEUROD1. Statistical significance was determined using a two-tailed Fisher's exact test followed by Benjamini–Hochberg correction.
E. Top enriched biological processes for upregulated DE genes after CRISPRa-NEUROG2. Statistical significance was determined using a two-tailed Fisher's exact test followed by Benjamini–Hochberg correction.

Figure S2: Details of TFome screen and hit transcription factors. Related to Figure 2.

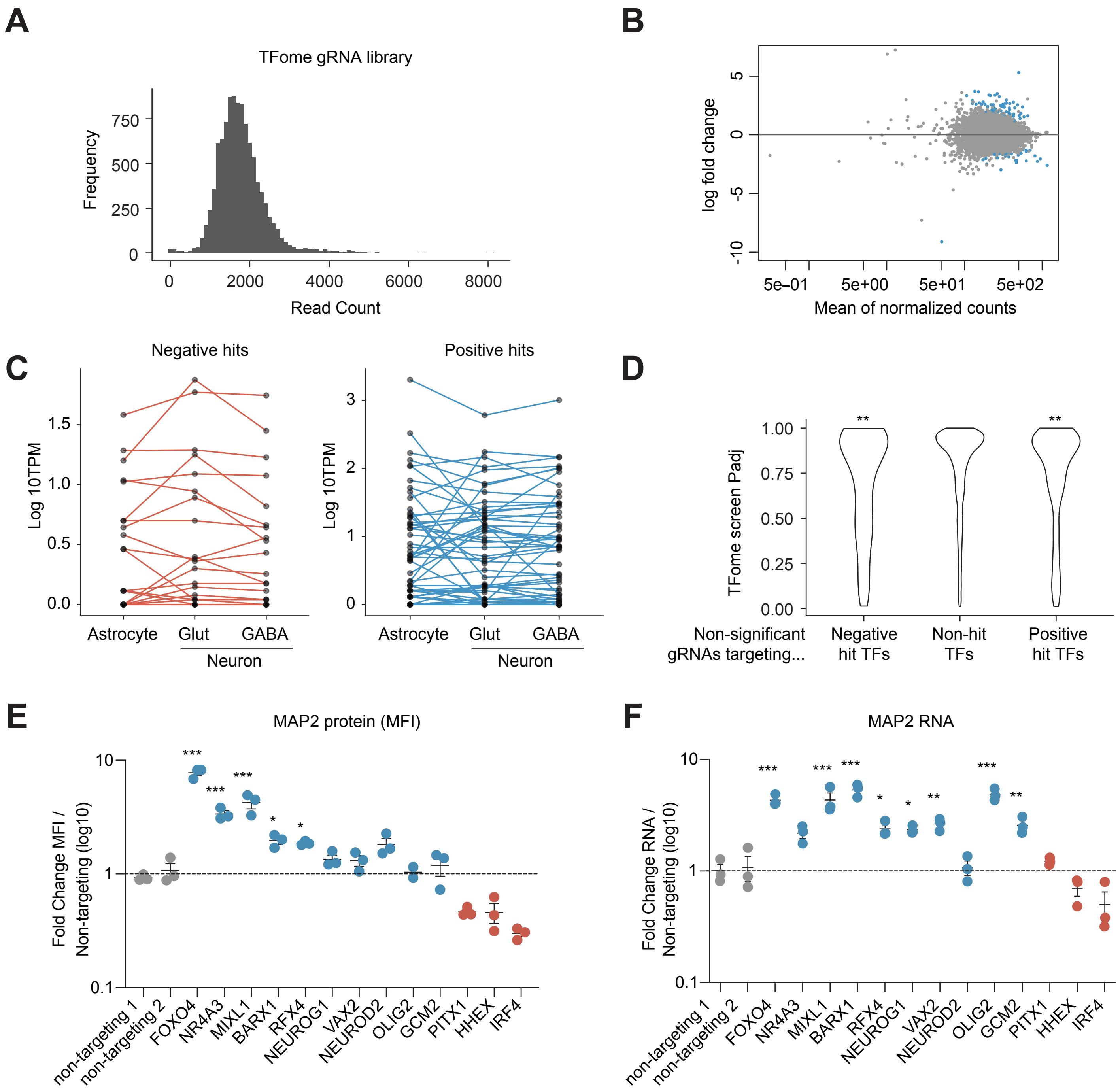

**Figure S2. Details of TFome screen and hit transcription factors. Related to Figure 2:**

A. Histogram of counts of gRNA protospacers in the TFome library plasmid pool.
B. Differential expression analysis of normalized gRNA counts between the MAP2-High and MAP2-Low cell populations. Blue data points indicate padj < 0.01 by differential DESeq2 analysis
C. Expression level of hit TFs in astrocytes or neurons in the Human Protein Atlas Single Cell Type data.
D. Violin plots of the distributions of adjusted p values of gRNAs targeting TFs that were negative hits, positive hits, or targeting non-hit TFs. **$p < 0.01$ by Mann-Whitney U test comparing distributions of negative or positive hit gRNAs to all gRNAs targeting non-hit TFs.
E. MFI of validations of selected hit factors for MAP2 protein expression 10 days post-transduction. **$p < 0.01$, ***$p < 0.001$ by global one-way ANOVA with Dunnett's post hoc test comparing all groups to non-targeting 1. Error bars represent SEM.
F. Validations of selected hit factors for MAP2 RNA levels 10 days post-transduction. *$p < 0.05$, ***$p < 0.001$ by global one-way ANOVA with Dunnett's post hoc test comparing all groups to non-targeting 1. Error bars represent SEM.

Figure S3: Cell-level analysis of astrocyte-to-neuron reprogramming perturb-seq screen. Related to Figure 3.

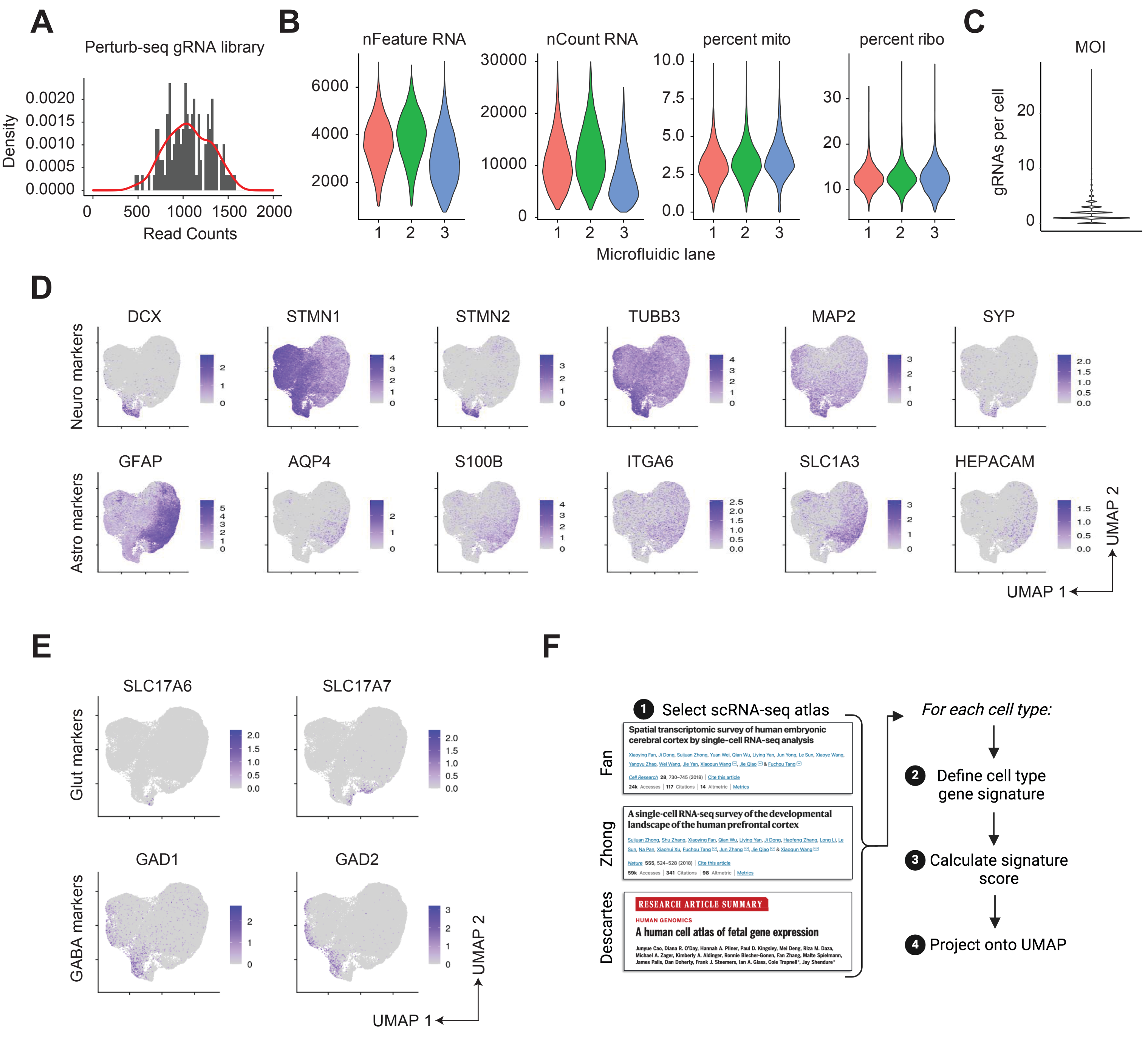

**Figure S3. Cell-level analysis of astrocyte-to-neuron reprogramming perturb-seq screen. Related to Figure 3:**

A. Histogram of counts of gRNA protospacers in the hit sublibrary plasmid pool.
B. Cell QC metrics for each microfluidic lane of the perturb-seq dataset.
C. Violin plot of the number of gRNAs detected in each cell after cell-gRNA link assignment.
D. Feature plots displaying count distribution of individual neuron (top) and astrocyte (bottom) marker genes in UMAP distribution.
E. Feature plots displaying count distribution of glutamatergic neuron (top) and GABAergic neuron (bottom) marker genes in UMAP distribution.
F. Schematic of the scRNA-seq atlas gene signature score assignment workflow.

Figure S4: CRISPRa perturbation potency and representation in perturb-seq screen.
Related to Figure 3.

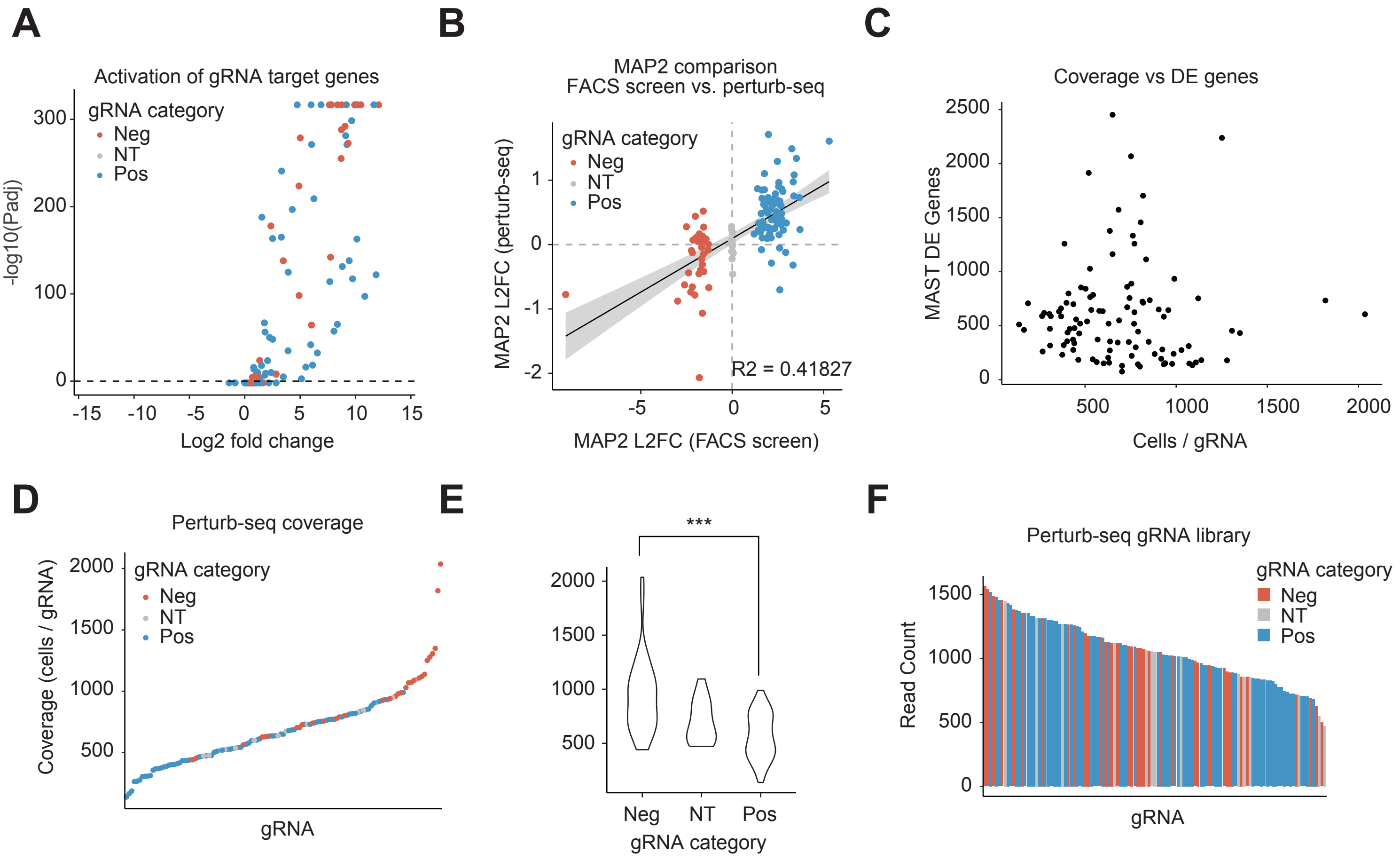

**Figure S4. CRISPRa perturbation potency and representation in perturb-seq screen. Related to Figure 3:**

A. Significance (Padj) versus fold change of CRISPRa TF-encoding target genes in scRNA-seq data after MAST.
B. Scatter plot comparing the fold change of MAP2 in the TFome screen vs. scRNA-seq.
C. Scatter plot comparing the coverage (number of cells with gRNA) and the number of DE genes ($p < .05$) called by MAST.
D. Ordered rank plot of coverage of each gRNA, colored by hit class.
E. Violin plot of distributions of coverages of gRNAs, separated by class. ***$p < 0.001$ by Mann-Whitney U test comparing negative and positive hit gRNAs.
F. Representation of gRNAs in the plasmid pool by hit class. Positive hits were not more lowly represented than other classes.

Figure S5: Grouping CRISPRa perturbations by shared transcriptome. Related to Figure 4.

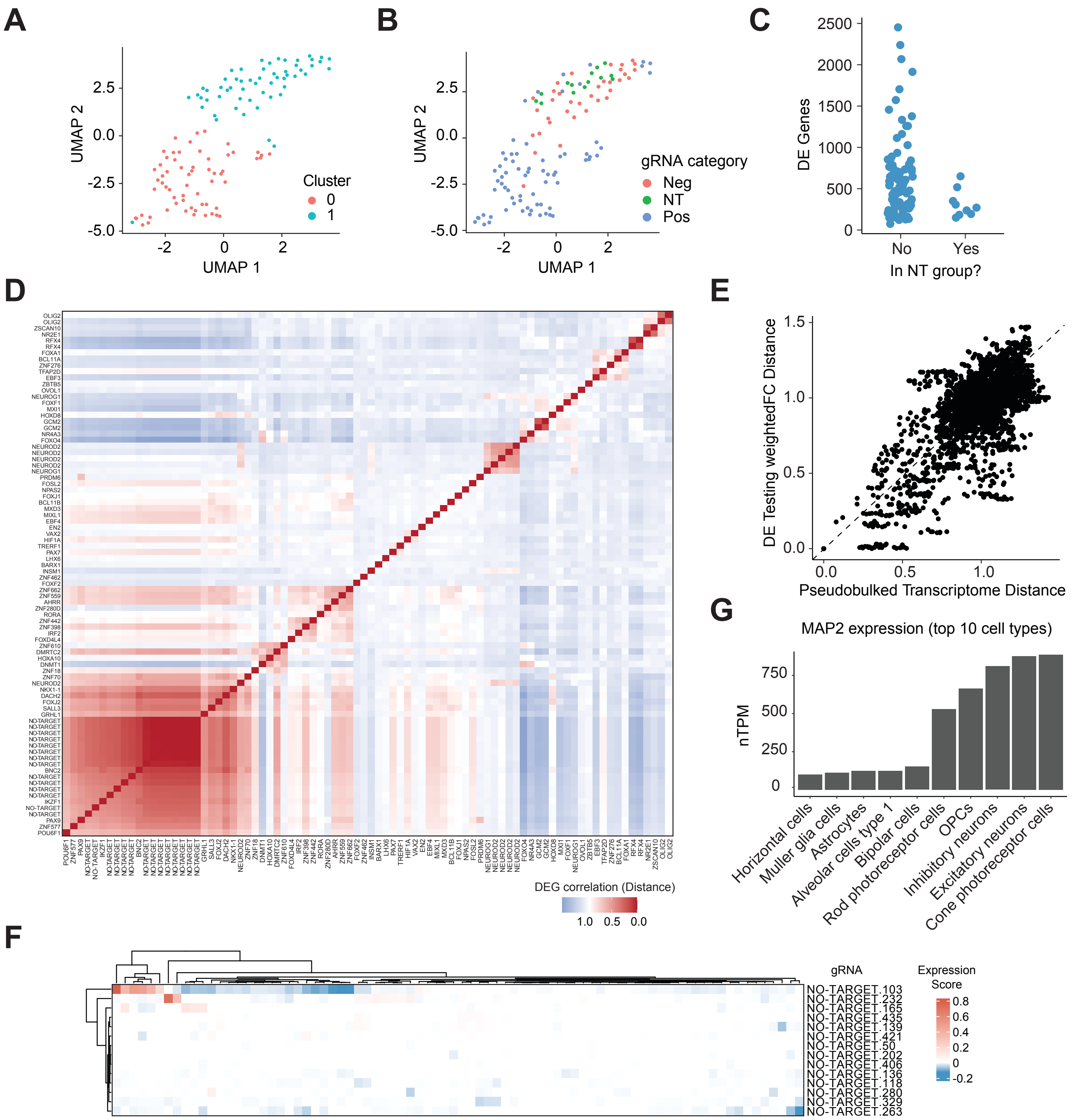

**Figure S5. Grouping CRISPRa perturbations by shared transcriptome. Related to Figure 4:**

A. Unsupervised clustering of pseudobulked perturbations, colored by assigned cluster.
B. Unsupervised clustering of pseudobulked perturbations, colored by hit class.
C. Number of DE genes of perturbations which group or do not group with non-targeting perturbations (related to Figure 4A).
D. Correlation of perturbations by weighted FC of DE genes. Positive or non-targeting gRNAs shown. Weighted FC = -log10(padj) * abs(fold change).
E. Scatter plot showing correlation of pairwise distances calculated by pseudobulked transcriptome (Figure 4A) and DE gene weighted FC (Figure S5D).
F. Atlas cell type gene signature module scores for non-targeting gRNAs. Module score values for all annotations are < 1.0.
G. Expression level of MAP2 in the top 10 cell types in the Human Protein Atlas Single Cell Type data.

Figure S6: TF candidates generate neuron-like cells from multiple systems and cell types.
Related to Figure 5.

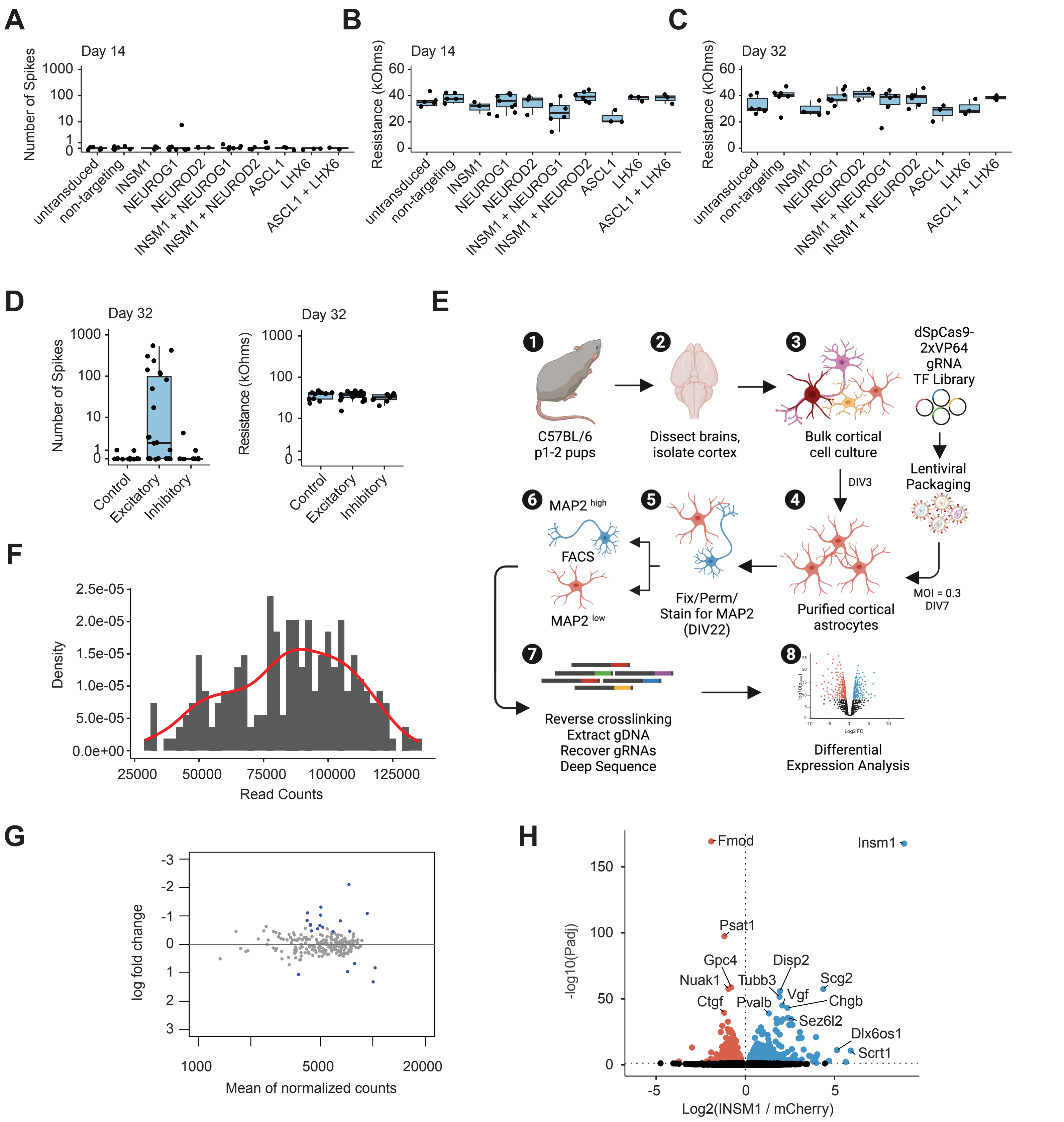

**Figure S6. TF candidates generate neuron-like cells from multiple systems and cell types. Related to Figure 5:**

A. Number of spikes (neuronal firing events) in 5-minute multi-electrode array recording 14 days post-transduction.
B. Resistance (kOhms) measured for each well in multi-electrode array recording 14 days post-transduction.
C. Resistance (kOhms) measured for each well in multi-electrode array recording 32 days post-transduction.
D. Number of spikes (neuronal firing events) and resistance (kOhms) in 5-minute multi-electrode array recording 32 days post-transduction, aggregated based on putative neuronal subtype.
E. Schematic of CRISPRa screening of mouse primary cortical astrocytes.
F. Histogram of counts of gRNA protospacers in the mouse screen sublibrary plasmid pool.
G. Differential expression analysis of normalized gRNA counts in the mouse screen between the MAP2-High and MAP2-Low cell populations. Blue data points indicate padj < 0.01 by differential DESeq2 analysis.
H. Significance (Padj) versus fold change in normalized RNA counts after RNA sequencing of INSM1 ORF overexpression vs. mCherry overexpression, 10 days post-transduction. DE genes are determined using a Wald test by DESeq2 (padj <0.01).

Figure S7: Details of paired screens for astrocyte-to-neuron reprogramming cofactors. Related to Figure 6.

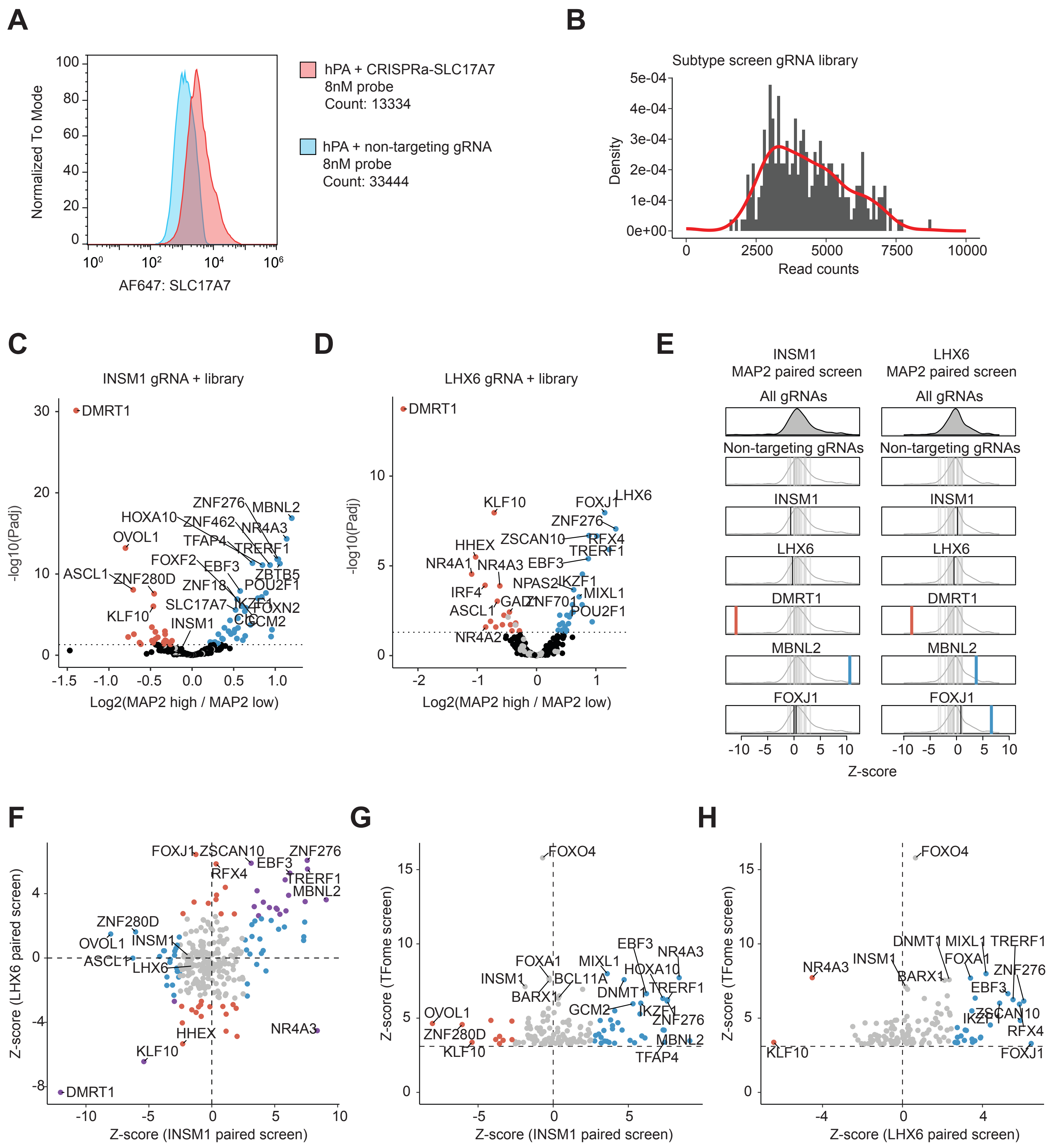

**Figure S7. Details of paired screens for astrocyte-to-neuron reprogramming cofactors. Related to Figure 6:**

A. Validation of SLC17A7 HCR-FlowFISH probeset for detection of intracellular RNA by flow-cytometry.
B. Histogram of counts of gRNA protospacers in the subtype screen library plasmid pool.
C. Significance (Padj) versus fold change in gRNA abundance between MAP2 high and MAP2 low populations in INSM1 paired CRISPRa screen.
D. Significance (Padj) versus fold change in gRNA abundance between MAP2 high and MAP2 low populations in LHX6 paired CRISPRa screen.
E. Z-scores of gRNAs for selected genes in INSM1 paired MAP2 (left) and LHX6 paired MAP2 (right) screens. Enriched gRNAs shown in blue and red (Padj < 0.01) were defined using a paired two-tailed DESeq2 test with Benjamini–Hochberg correction.
F. Scatter plot of z-score of gRNA abundance in the INSM1 MAP2 paired screen and the LHX6 MAP2 paired screen.
G. Scatter plot of z-score of gRNA abundance in the INSM1 MAP2 paired screen and the TFome single-factor MAP2 screen. gRNAs present in both screens are shown.
H. Scatter plot of z-score of gRNA abundance in the LHX6 MAP2 paired screen and the TFome single-factor MAP2 screen. gRNAs present in both screens are shown.

Figure S8: Chromatin changes after paired CRISPRa reprogramming. Related to Figure 6.

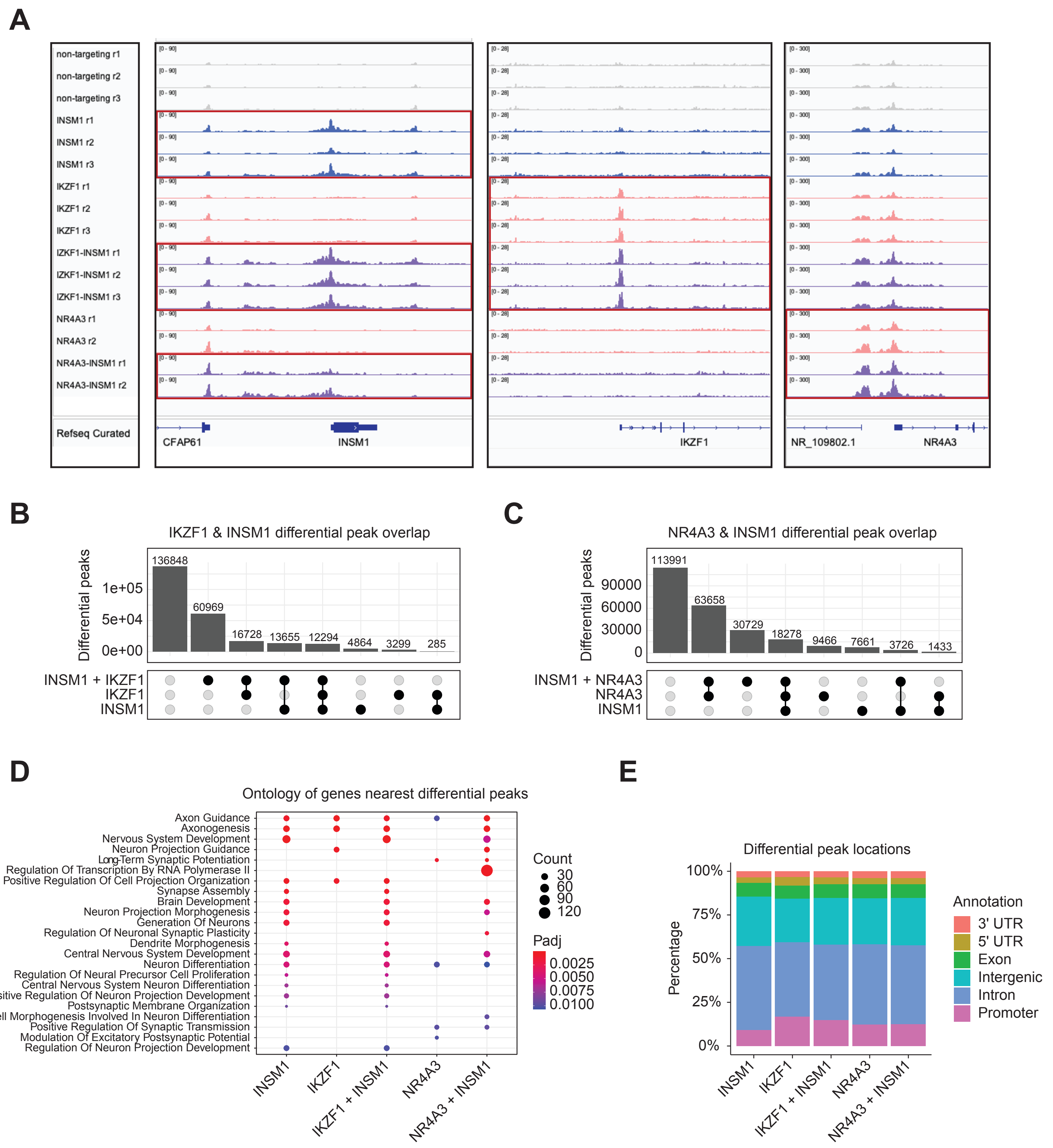

**Figure S8. Chromatin changes after paired CRISPRa reprogramming. Related to Figure 6:**

A. Browser tracks of ATAC-seq (reads per kilobase per million mapped reads [RPKM]-normalized BigWig, bin size = 25bp) at loci of targeted TFs. Conditions where the displayed TF are directly targeted by a delivered gRNA are shown in red boxes.
B. Upset plots of IKZF1 & INSM1 and combination differentially accessible peaks. Differential peaks (padj <.01) for each sample were determined by DESeq2 vs. non-targeting gRNA.
C. Upset plots of NR4A3 & INSM1 and combination differentially accessible peaks.
D. Top enriched biological processes for genes nearest top 1000 differentially accessible peaks by z-score after perturbation of single factors or combinations. Statistical significance of term enrichment was determined using a two-tailed Fisher's exact test followed by Benjamini–Hochberg correction.
E. Stacked bar of differentially accessible peaks by genomic element type.
